# Dynamic *Phaeodactylum tricornutum* Exometabolites Shape Surrounding Bacterial Communities

**DOI:** 10.1101/2022.06.08.495228

**Authors:** Vanessa Brisson, Courtney Swink, Jeffrey Kimbrel, Xavier Mayali, Ty Samo, Suzanne M. Kosina, Michael Thelen, Trent R. Northen, Rhona K. Stuart

## Abstract

The roles of exometabolites in mediating algal-bacterial interactions and regulating microbial community composition are not well understood. Here, we identified specific exometabolites from the model diatom *Phaeodactylum tricornutum* affecting abundance of specific bacterial taxa in isolation and in a community setting. We examined the response of a *P. tricornutum*-adapted enrichment community and found that both algal exudates and algal presence drove similar changes in community composition compared to controls. Using LC-MS/MS, we identified 50 metabolites produced by axenic *P. tricornutum* and found that different exometabolites accumulated during different algal growth phases. Profiling growth of 12 bacterial isolates representative of the enrichment community uncovered two algal exometabolites (out of 12 tested) which supported growth of a subset of isolates as a primary carbon source. We compared enrichment community response with and without the addition of two contrasting metabolites: 4-hydroxybenzoic acid, which supported isolate growth, and lumichrome, which did not. Exogenous metabolite additions did promote increased abundances of taxa that were able to utilize the metabolite in the isolate study, but also revealed the importance of factors relating to algal presence in regulating community composition. Collectively, this work demonstrates the influence of specific algal exometabolites in driving microbial community composition.

## INTRODUCTION

Microalgae play important roles in global carbon cycling and industrial applications for bioproduct and biofuel production (1–3). As with land plants and other host-microbial systems, microalgal activity, productivity, and stability are closely tied to surrounding microbial communities (4). As primary producers, algae provide fixed carbon that supports microbial growth (5). Bacteria, in turn, can have both beneficial and detrimental effects on their algal hosts. Beneficial bacteria produce necessary vitamins and growth promoting compounds and provide protection from pathogens, while harmful bacteria produce detrimental effects through competition and pathogenesis (5, 6). In some cases, bacteria switch from growth promotion to algicidal activity as their algal hosts age (7, 8). Regardless of whether interactions are helpful or deleterious, a more detailed understanding of the chemical ecology driving these relationships will inform our predictive understanding of these systems.

Exometabolite release is hypothesized to be a primary mechanism by which host organisms influence microbiome composition. Previous work showed that different algal hosts assemble different phycosphere communities (15) and different algae have distinct exometabolomes (11), highlighting the potential importance of individual algal exometabolomes or specific exometabolites in governing algal-bacterial interactions. Further, microbial abundances have been shown to change in response to metabolite composition in simplified systems with defined suites of metabolites (16). The potential for exometabolome composition to influence specific bacterial taxa is also suggested by the selective use of different exometabolites by bacterial isolates in co-culture with the diatom *Thalassiosira pseudonona* (12). Bacterial growth and motility, both factors that can modulate the microbial community, were influenced by metabolites (rosmarinic and azelaic acids) detected in exudates from the diatom *Asterionellopsis glacialis* (14). P-coumarate, a metabolite released by *Emiliana huxleyi* during senescence, has been linked to a shift from mutualistic to algicidal activity of the bacterium *Phaeobacter inhibens* (17). Given these impacts of exometabolites on algal-bacterial interactions, characterization of algal exometabolomes and their roles is critical for understanding these systems.

Intracellular metabolites have been used as a proxy for exometabolites because they become available outside the cell through diffusion, exudation, and cell lysis (4, 9). Used in conjunction with transcriptomics, this approach has produced insights into algal-bacterial interactions and metabolite exchange dynamics (10). However, recent work with several algal strains suggests that the algal exometabolome composition differs significantly from the endometabolome (11). Thus, direct analysis of the exometabolome is needed to further understand algal bacterial interactions. Algal exometabolomics, particularly for saltwater algae, presents technical challenges due to high salts and low metabolite concentrations (12), but recent studies have started to address this knowledge gap. The exometabolomes of the eukaryotic microalgae *Asterionellopsis glacialis*, *Thalassiosira pseudonona*, *Thalassiosira rotula*, *Phaeodactylum tricornutum*, *Microchloropsis salina*, *Chlamydomonas reinhardtii*, and *Desmodesmus intermedius* have been characterized using untargeted and targeted metabolomic approaches (11–14). As exometabolite characterization expands, the next challenge is to integrate that knowledge with the effects of individual bacterial strain responses to understand interactions in a complex microbiome.

In this study, we investigate the role of exometabolites from the model diatom *Phaeodactylum tricornutum* in shaping surrounding microbial communities. We set out to test the hypothesis that algal exudates drive microbial community composition and to identify specific algal exometabolites that can be linked to specific changes in algal-associated microbial communities. We used microbial community analysis to investigate responses to algal exudates and algal presence using a previously described microbial community enriched for growth with *P. tricornutum* (15, 18). We conducted a time resolved metabolomic analysis to characterize exudates from axenic *P. tricornutum* and identify specific exometabolites of interest. Using selected identified metabolites, we conducted growth assays with bacterial isolates previously characterized for their associations with *P. tricornutum* (19, 20). Based on results from these assays, we again used microbial community analysis to investigate the role of specific identified exometabolites in the context of a complex microbial community with and without algae and algal exudates, allowing us to investigate the interacting roles of exometabolites, algae, and bacteria.

## MATERIALS AND METHODS

### Organisms

We used the model diatom *P. tricornutum* strain CCMP 2561. Bacterial isolates were isolated in our lab and have been described elsewhere (19, 20). Information on their origins and identifications is summarized in Table 1. Enrichment cultures were from the same phycosphere enrichments used to obtain the bacterial isolates and have been described elsewhere (15, 18).

**Table 1.** Bacterial isolates and corresponding microbial community ASVs with exact matches to the 16S-V4 region. Isolate origin and identification are from Mayali et al. 2022 (19).

| Isolate Origin | Bacterial Isolate | Exact | Mean Relative Abundance |  |  |
| --- | --- | --- | --- | --- | --- |
|  |  | Match | Under Different Growth Conditions |  |  |
|  |  | 16S-V4 | *P. tricornutum* | *P. tricornutum* | Algal-Free |
|  |  | ASV | Present | Spent Medium | Control |
| phycosphere enrichments | *Oceanicaulis 13A* | ASV12 | 8.1% | 4.4% | 0.3% |
|  | *Roseibium 13C1* | ASV6 | 2.5% | 6.8% | 3.5% |
|  | *Thalassospira 13M1* | ASV23 | <0.1% | 0.5% | 0.1% |
|  | *Yoonia 4BL* | - | - | - | - |
|  | *Arenibacter ARW7G5Y1* | ASV34 | <0.1% | 0.5% | <0.1% |
|  | *Algoriphagus ARW1R1* | ASV9 | 7.7% | 8.5% | 0.1% |
|  | *Stappia ARW1T* | - | - | - | - |
|  | *Devosia EAB7W2* | ASV18 | 0.2% | 2.6% | <0.1% |
|  | *Alcanivorax EA2* | ASV14 | 0.1% | <0.1% | 12.3% |
|  | *Marinobacter 3-2* | ASV1 | 12.2% | 33.5% | 30.2% |
| seawater | *Amylibacter N2S* | - | - | - | - |
|  | *Sulfitobacter N5S* | - | - | - | - |
| **Total** |  |  | **30.9%** | **56.9%** | **46.5%** |

### Algal exometabolomics

Prior to the experiment, 125 mL glass Erlenmeyer flasks were baked at 500 °C for two hours to remove residual organic matter. Axenic algae were grown in 50 mL Enriched Saltwater Artificial Water (ESAW) medium (11, 21–23). Flasks were incubated with shaking (90 rpm, 22 °C, 12-hour day/night cycle, 3500 lux daytime illumination). Twenty replicate flasks were inoculated with two mL each from a one-week-old *P. tricornutum* culture. Algal growth was monitored by measuring chlorophyll A fluorescence using a Trilogy Fluorometer [Turner Designs, San Jose, CA, USA] with the Chlorophyll In-Vivo Module. At each metabolite collection time point (0, 3, 6, 9, and 12 days) (Supplementary Figure S1), four flasks were destructively sampled for metabolomics analysis of spent medium and cell pellets as previously described (11).

For solid phase metabolite extraction, samples (filtered spend medium and filtered water extracts of disrupted cell pellets) were thawed, and 30 mL of each sample was acidified by adding 300 µL of 1 M HCl. Bond Elut PPL columns [Agilent, Santa Clara, CA, USA] were rinsed with 3 mL HPLC grade methanol followed by 6 mL ultrapure water. Acidified samples were passed through prepared columns. Columns were rinsed with 3 mL of 0.01 M HCl and allowed to air dry for 5 minutes. Metabolites were eluted with 1 mL HPLC grade methanol, dried in a vacuum centrifuge, and resuspended and prepared for analysis as previously described (11, 24, 25).

Metabolomes were analyzed with liquid chromatography tandem mass spectrometry (LC-MS/MS). LC-MS/MS instrumentation and parameters are detailed in Supplementary Table S1. Metabolite identifications were obtained using Metabolite Atlas (https://github.com/biorack/metatlas) to identify features with *m/z*, retention time, and fragmentation spectra matched to a library of analytical standards run in our lab under the same LC-MS/MS analysis conditions (26, 27).

### Bacterial isolate growth assays

Twelve selected exometabolites were tested for their ability to support growth as the primary carbon source for each of the bacterial isolates. For each metabolite, ESAW medium was prepared with the added metabolite at a total carbon concentration of 60 ppm. This concentration was chosen as approximately ten times the previously observed DOC concentration in *P. tricornutum* spent medium (11) in order to stimulate sufficient growth for detection. Each metabolite was tested with each bacterial isolate in triplicate wells with replicate positions randomized across the 96-well flat-bottom plates. Negative controls (no added metabolite) were also included. Plates were incubated with shaking (90 rpm, 24 °C). OD 600 was measured daily for 8 days using a Cytation5 plate reader [BioTek, Winooski, VT, USA].

### Microbial community response

Three algal conditions were tested: algae present, algal spent medium, and algal-free control. Experiments were conducted in flasks as described above for the metabolomics experiment. For the algae present condition, flasks were prepared with ESAW and inoculated with 2 mL of one-week-old *P. tricornutum* culture at the same time as bacterial inoculation. For the algal spent medium condition, flasks were prepared with spent medium from a one-week-old *P. tricornutum* culture that was filtered through a 0.2 micron filter to remove algal cells. Phosphate and nitrate were replenished in algal spent medium prior to inoculation with bacteria. Algal-free control flasks were prepared with ESAW medium without addition of algae, algal spent medium, or other carbon source beyond what was present in the base medium.

For added metabolite experiments, the same three algal conditions as above were prepared with the addition of 0.01 mM of the metabolite being tested. This concentration was selected (A) so that the added metabolite represented a small change and did not overwhelm algal metabolites present in the algal spent medium and algae present conditions, and (B) because previous work with lumichrome indicated that this concentration impacted *P. tricornutum* growth (11).

To obtain bacterial inoculum, two previously described phycosphere enrichment cultures were combined (15, 18). Enrichments were grown in ESAW medium for one week. Cultures were filtered (0.8 µm pore size, 45 mm diameter Supor membrane filter [Pall Corporation, Port Washington, NY, USA]) to remove algal cells but allow bacterial cells to pass through. Filtrate was used to inoculate all flasks with 2 mL (1 mL from each enrichment).

Twenty-five mL samples were collected from each flask after 6, 8, and 10 days of incubation. Flasks were replenished with 25 mL of the appropriate fresh or algal spent medium. One mL from each sample was fixed for flow cytometry by adding 100 µL 37% formaldehyde. The remaining sample was filtered onto a 45 mm diameter, 0.2 µm pore size Supor membrane filter [Pall Corporation, Port Washington, NY, USA]. Filters were frozen at −80 °C until DNA extraction. DNA extractions were conducted using the NucleoSpin 96 Tissue Kit [Macherey-Nagel, Düren, Germany] with modifications to the vacuum processing protocol as detailed in the Supplemental Methods. DNA concentrations were quantified using the Invitrogen Qubit dsDNA HS assay kit [Thermo Fischer Scientific, Waltham, MA, USA] and stored at −20 °C until sequencing.

Sequencing was performed through Laragen, Inc. [Culver City, CA, USA], using the earth microbiome project protocol. The 515F (GTGYCAGCMGCCGCGGTAA) (28) and 806R (GGACTACNVGGGTWTCTAAT) (29) primers were used to amplify and sequence the 16S-V4 region. Sequencing was performed on an Illumina MiSeq [Illumina, San Diego, CA, USA] using the MiSeq V2 PE150 reagent kit. Adapter sequences were trimmed from reads using cutadapt, and trimmed reads were processed using the DADA2 package (version 1.20.0) to assess read quality, filter and trim reads, merge paired reads, remove chimeras, and generate an amplicon sequence variant (ASV) table (30). ASV taxonomy was assigned using both the RDP database (version 18) and the Silva database (version 138). Where identifications conflicted, the RDP identification is given with the Silva identification in parentheses. Microbial community analysis was conducted using phyloseq (version 1.36.0) (31). Chloroplast and mitochondrial sequences were removed, as were ASVs that were not detected with at least ten counts in at least four samples. The ANCOM-BC package (version 1.2.2) was used to assess differential abundance and to generate bias corrected ASV counts (32).

## RESULTS

### *P. tricornutum* exudates and algal presence drive microbial community composition

We investigated the impacts of algal presence and algal exudates on a microbial community that had previously been enriched to grow with algae (15, 18). Microbial community analysis revealed this to be a relatively simplified bacterial community. After removing mitochondrial and chloroplast reads and low prevalence ASVs (present in fewer than four samples with at least ten reads), there were 62 bacterial ASVs and no archaeal ASVs. All 62 ASVs could be identified to at least the family level, and 58 could be identified at the genus level, representing 42 genera. Half of the ASVs were low abundance (< 0.1% mean relative abundance), while five were high abundance ASVs (> 5% mean relative abundance) (Supplementary Figure S2).

The composition of microbial communities grown with algal spent medium from late-log (day 7) or with algae present were distinct from algal-free controls and from each other (Figure 1, Supplementary Figure S2). In the principal coordinate analysis, microbial communities grown with algal spent medium were intermediate between those grown with algae present and algal-free controls along the first principal coordinate axis, which explained 55.3% of the variance (Figure 1). The second principal coordinate axis further differentiated communities grown on algal spent medium from the other conditions and differentiated between time points within each condition. Permutational analysis of variance (PERMANOVA) indicated that algal condition (algae present, algal spent medium, or control), time, and their interaction were all significant drivers of microbial community composition (Supplementary Table S2). Community alpha diversity, as measured by Shannon Diversity Index, was driven by algal presence, and also increased over time (Supplementary Table S3, Supplementary Figure S3).

**Figure 1.**
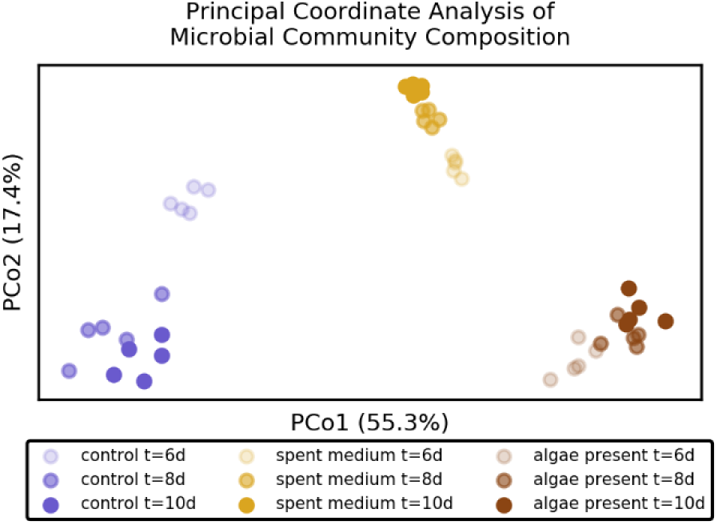
Principal coordinate analysis showing differences in microbial community composition using Jensen-Shannon Divergence between samples counts. Each point represents one sample. Colors represend different growth conditions of algae present (brown), algal spent medium (yellow), or algal-free control (blue). Shading indicates the time of sample collection at 6 days (light), 8 days (medium), or 10 days (dark).

Algal presence and spent medium shifted relative abundances of most ASVs in the same direction compared to controls (Figure 2, Supplementary Figure S2). Of the 31 ASVs with mean relative abundance of at least 0.1%, 27 were significantly differentially abundant between algal spent medium and controls, and a distinct set of 28 were differentially abundant between algae present and controls (Figure 2-a, Supplementary Figure S4). Ten ASVs increased in relative abundance with both algae and algal spent medium compared to controls, and nine decreased with both algae and algal spent medium compared to controls. Only six had divergent responses to algal presence and spent medium. These included four (*Marinobacter* ASV3, *Roseibium* (*Labrenzia)* ASV6, *Thalassospira* ASV7, and *Thalassospira* ASV23) which increased with spent medium but decreased with algae, and two (*Roseitalea* ASV 29 and *Maricaulis* ASV32) which decreased with spent medium but increased with algae. Regression analysis indicated that there was a significant correlation (P < 0.001, R^2^ = 0.51) between the responses of ASVs (log2 fold changes compared to control) to algal spent medium and to algal presence (Figure 2-b).

**Figure 2.**
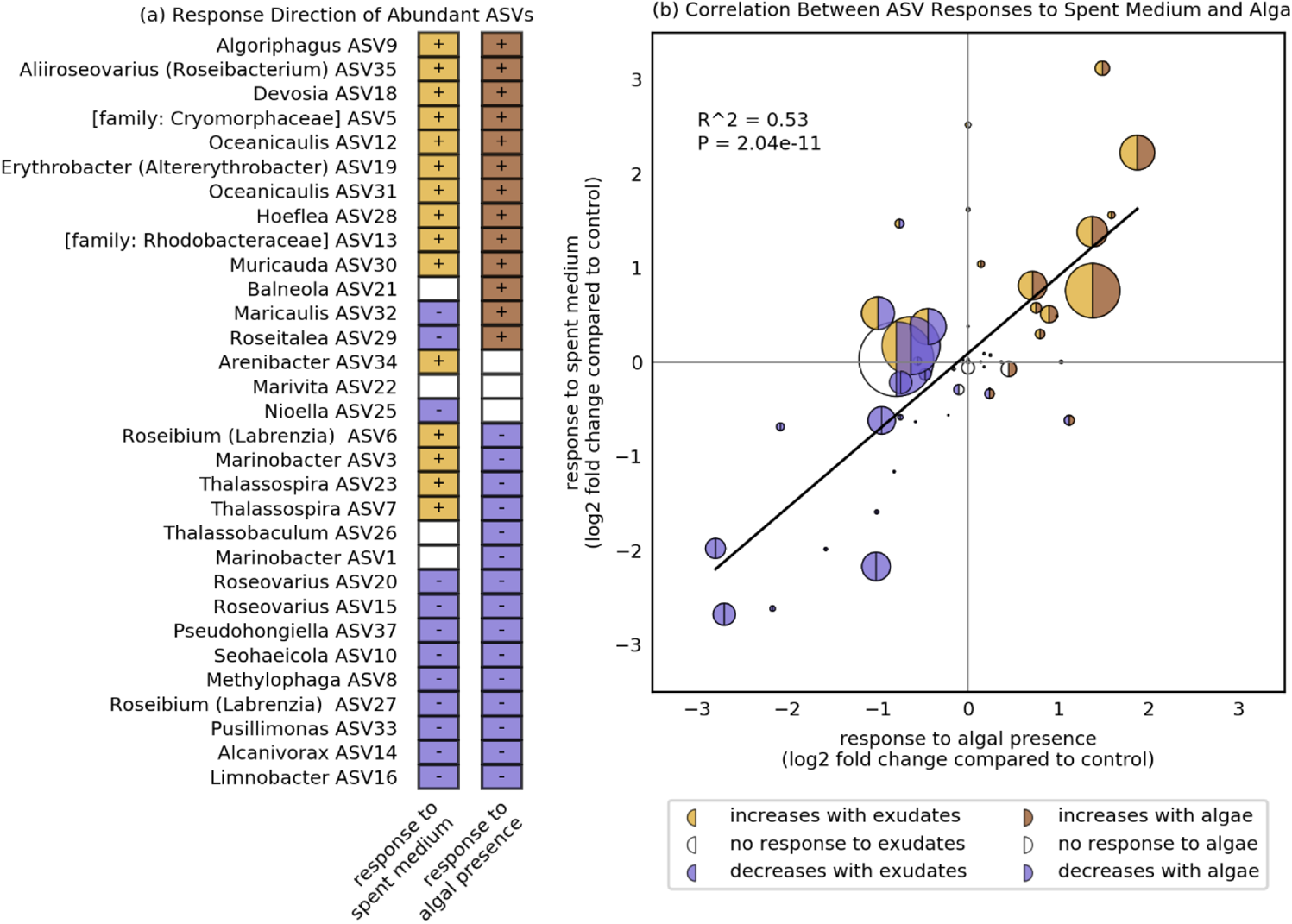
Response of individual ASVs to algal presence and algal spent medium compared to algal-free controls. (a) Direction of ASV responses to algal presence and algal spent medium. Only the ASVs with at least 0.1% mean relative abundance are included. “+” indicates a statistically significant increase in relative abundance compared to conrol, and “-” indicates a significant decrease compared to control. (b) Correlation between ASV responses to spent medium and algal presence. Each circle represents one ASV. Circle size is proportional to mean relative abundance. Shading indicates statistically significant responses to algal presence (left side of circle) and algal spend medium (right side of circle). All 62 ASVs are included in this plot. Diagonal line represents the best fit linear regression.

### *P. tricornutum* produces diverse exometabolites

We identified 50 metabolites produced by *P. tricornutum*, including organic acids, vitamins, amino acids, and nucleosides/nucleobases, as well as derivatives of these (Figure 3, Supplementary Tables S4 and S5, Supplementary Figure S5). The majority of identified metabolites (43 out of 50) were nitrogen containing compounds. Metabolite profiles from cell pellets and spent medium were distinct from each other, and most individual metabolites were highly associated with one or the other sample type (Supplementary Figure S5). Based on hierarchical clustering, 26 metabolites were identified as primarily exometabolites and 24 were identified as primarily cell pellet metabolites. As a compound class, organic acids were more highly associated with the spent medium (Chi-squared test p-value = 0.002). Although we observed trends of higher associations of nucleosides/nucleobases and amino acids with cell pellets and higher association of vitamins with spent medium, these associations were not statistically significant. Three indole containing metabolites (indole3-acetic acid, indole-3-pyruvic acid, and 5-hydroxyindoleacetic acid) and the additional plant hormone salicylic acid were detected, and were all significantly more abundant in the spent medium than in the cell pellets.

**Figure 3.**
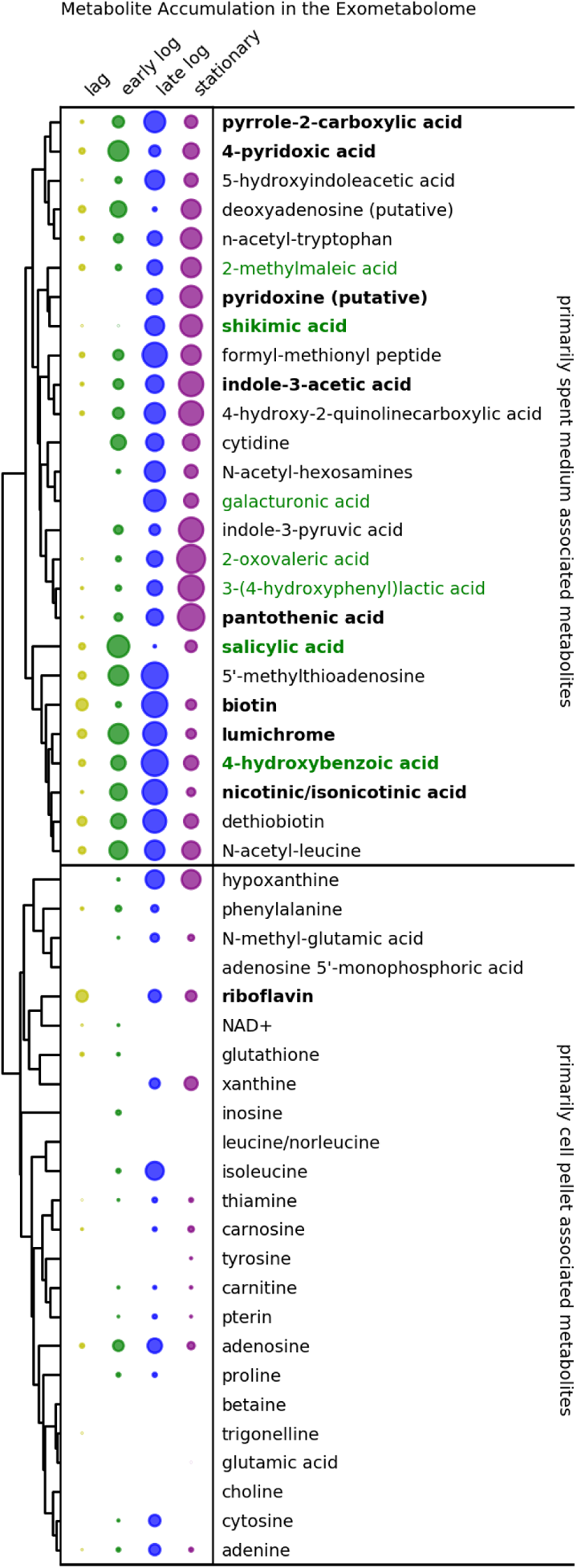
Exometabolite accumulation during different growth phases. Circle size represents the mean accumulation (relative intensity increase) during each growth phase. For metabolite levels in individual samples refer to Supplementary Figure S5. Growth phase colors correspond to colors in Supplementary Figure S1. Dendrogram to the left shows hierarchical clustering based on metabolite concentration profiles. Metabolite name colors indicate nitrogen content (black = contains nitrogen, green = does not contain nitrogen). Metabolite names in bold were used in the bacterial isolate growth experiment.

Exometabolite production was dynamic, with different metabolites accumulating during different algal growth phases (Figure 3). Hierarchical clustering grouped metabolites based on their accumulation patterns. Two exometabolites (galacturonic acid and pyridoxine) only accumulated during late log and stationary phases, but were not detected earlier in growth. At the other extreme, 5’-methylthioadenosine accumulated through log phase but did not continue to accumulate in stationary phase.

### Individual exometabolites impact bacterial isolate growth

Based on their accumulation in the exometabolome, 12 metabolites were selected to further test their impacts on the growth of a select set of 12 bacterial isolates. Six of the metabolites were vitamins or vitamin derivatives: pantothenic acid (vitamin B5), riboflavin (vitamin B2), biotin (vitamin B7), nicotinic acid (vitamin B3), lumichrome (degradation product of vitamin B2), and pyroxidine (vitamin B6). The remaining six metabolites were organic acids: pyrrole-2 carboxylic acid, 4-hydroxybenzoic acid, 4-pyridoxic acid, indoel-3-acetic acid, salicylic acid, and shikimic acid. The bacterial isolates were representative of ASVs found in the microbial communities. Of the ten bacterial isolates originating from phycosphere enrichments, eight had 100% sequence identity between their 16S sequence and an ASV sequence from the enrichment community experiment (Table 1). Together these exact match ASVs represented between 30% and 57% total relative abundance under the different algal growth conditions. No close matches were found to the 16S sequences from the two non-phycosphere bacterial isolates originating from seawater (*Amylibacter* NS2 and *Sulfitobacter* NS5) or to the other two phycosphere derived bacterial isolates (*Stappia* ARW1T and *Yoonia* 4BL).

A subset of exometabolites supported growth of specific bacterial isolates as the primary source of organic carbon (ESAW medium contains a small amount of organic carbon in vitamins and EDTA as described in the Supplemental Methods). Of the 12 metabolites tested, two supported growth, as measured by a significant increase in optical density, of at least one of the 12 bacterial isolates (Figure 4, Supplementary Figure S6). Both metabolites that supported growth were organic acids and not vitamins or their derivatives. 4-hydroxybenzoic acid was the most widely utilized substrate under the conditions tested and supported the growth of four isolates: *Roseibium* 13C1, *Thalassospira* 13M1, *Stappia* ARW1T, and *Alcanivorax* EA2. Shikimic acid supported the growth of *Stappia* ARW1T and *Alcanivorax* EA2.

**Figure 4.**
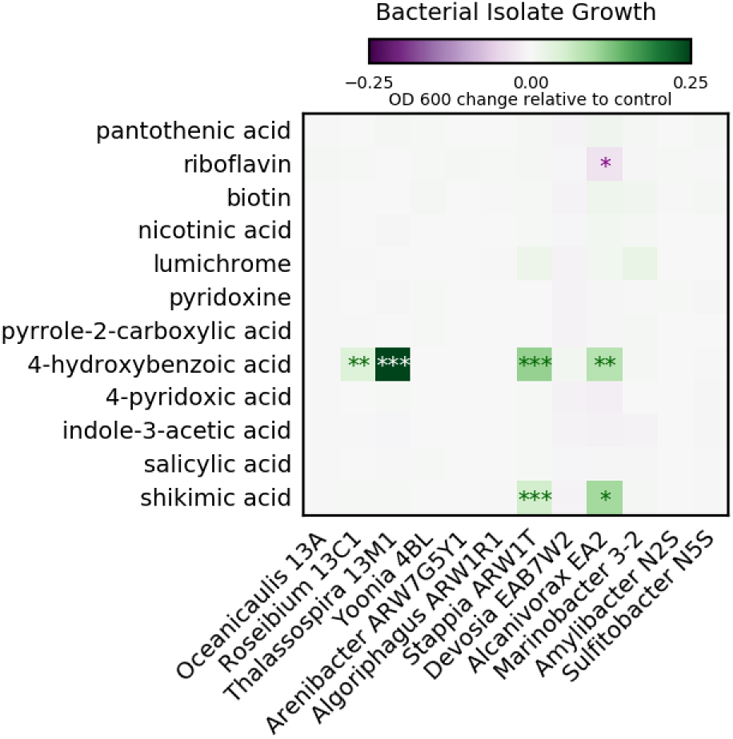
Growth of algal-associated bacterial isolates on select algal exometabolites. Heatmap color intensity corresponds to the mean increase (green) or decrease (purple) in final optical density for each isolate and metabolite combination compared to control for that isolate with no added metabolite. Means are for three biological replicates. Statistically significant differences compared to control with no added metabolite are indicated based on p-values with significance codes: 0 ‘***’ 0.001 ‘**’ 0.01 ‘*’ 0.05 ‘.’ 0.1 ‘ ’ 1. For individual datapoints contributing to the heatmap for the metabolites that supported growth, refer to Supplementary Figure S6.

To relate responses to 4-hydroxybenzoic acid to genetic potential, we searched the previously sequenced genomes of these isolates (19) for genes for five enzymes related to 4-hydroxybenzoic acid degradation: 4-hydroxybenzoate 3-monooxygenase (gene: *pobA*, EC:1.14.13.2), 4-hydroxybenzoate 1-hydroxylase (EC:1.14.13.64), 4-hydroxybenzoate decarboxylase (EC:4.1.1.61), 4-hydroxybenzoate polyprenyltransferase (gene: *ubiA*, EC:2.5.1.39), and 4-hydroxybenzoate-CoA ligase (EC:6.2.1.27). Of these only *pobA* and *ubiA* were identified in the isolate genomes. All isolate genomes except for *Algoriphagus* ARW1R1 and *Arenibacter* ARW7G5Y1 contained *ubiA*, while *pobA* was found in the genomes of seven of the twelve isolates (Supplementary Table S7), including three (*Thalassospira* 13M1, *Rosibium* 13C1, and *Stappia* ARW1T) which grew on 4-hydroxybenzoic acid.

### Specific exometabolites influence relative abundance of bacteria in a complex community

Two algal exometabolites, 4-hydroxybenzoic acid and lumichrome, were selected for further study of their impacts on bacterial community compositions. 4-hydroxybenzoic acid was selected as a potential bacterial growth substrate because it selectively supported the growth of specific bacterial isolates in the experiment above. Lumichrome was selected as a representative vitamin derivative because both this study and a previous study found significant production by *P. tricornutum*, and the previous study found that it influenced *P. tricornutum* growth as well (11). Both metabolites had similar accumulation profiles, with maximum accumulation during log phase growth (Figure 3). Because all microbial community experiments were conducted, sequenced, and analyzed together, this analysis contains the same 62 ASVs described above.

Exogenous additions of metabolites had significant impacts on bacterial community compositions, both with and without *P. tricornutum* present. As with the communities without metabolite addition, the first principal coordinate, which accounted for 30.9% of the variance, separated samples primarily by algal treatment (algae present, algal spent medium, and control), with algal spent medium being intermediate to algae present and algal-free controls (Figure 5). The second and third principal components, accounting for 11% and 9% of the variance respectively, separated samples by metabolite addition, especially separating the samples with added 4-hydroxybenzoic acid from those with lumichrome or no added metabolite. PERMANOVA indicated that algal condition, metabolite addition, time, and their interactions were all significant factors affecting microbial community composition (Supplementary Table S6).

**Figure 5.**
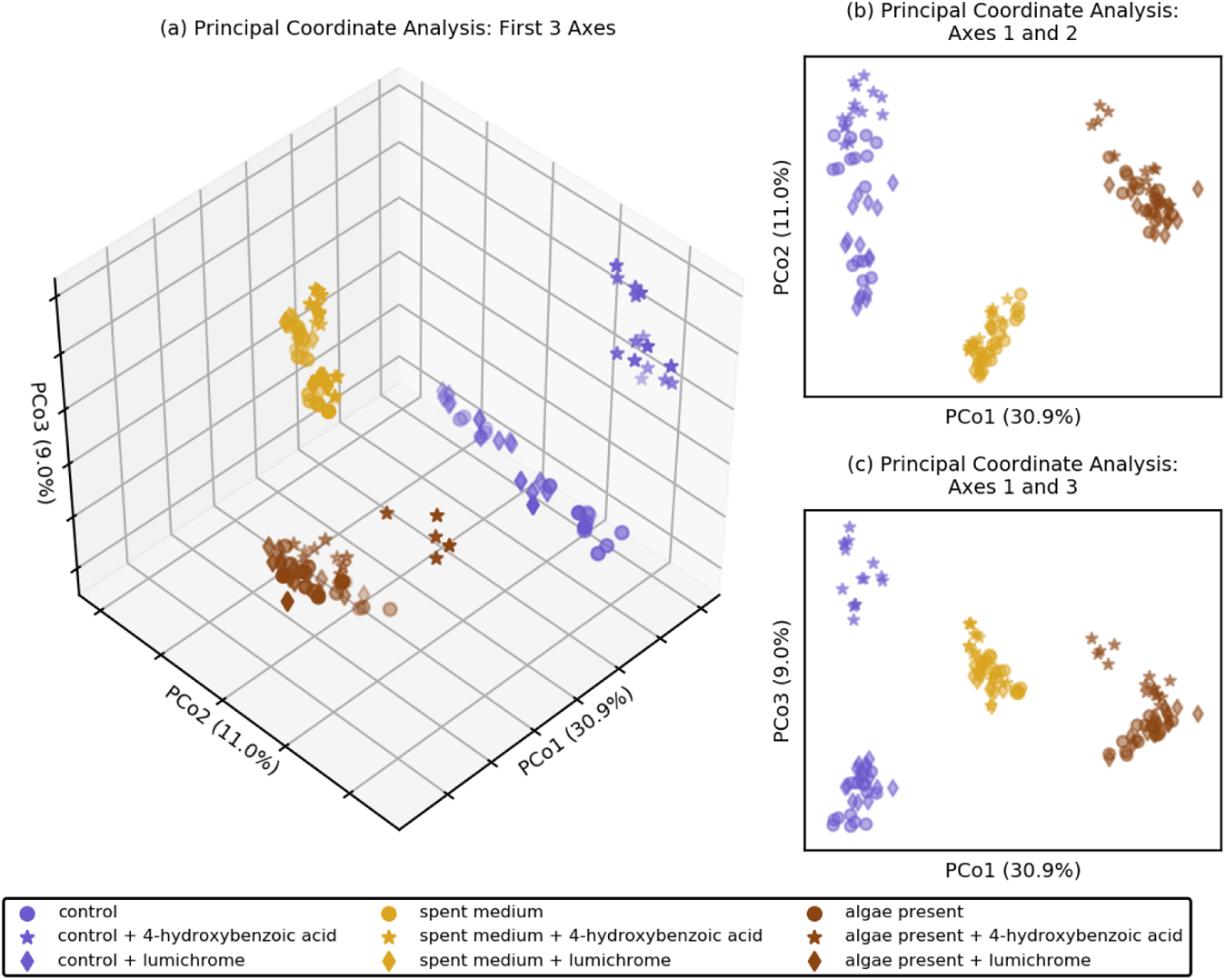
Principal coordinate analysis showing differences in microbial community composition in response of exogenous addition of select metabolites based on Jensen-Shannon Divergence between samples. (a) Three-dimensional plot of the first three principal components. (b) Principal coordinates 1 and 2. (c) Principal coordinates 1 and 3. Each point represents one sample. Colors represent different base growth conditions of algae present (brown), algal spent medium (yellow), or algal-free control (blue). Different shapes indicate different added metabolites: lumichrome (diamonds), 4-ydroxybenzoic acid (stars), and no metabolite addition controls (circles).

Addition of 4-hydroxybenzoic acid drove significant changes in the relative abundances of several ASVs. Both *Thalassospira* ASVs (ASV7 and ASV23), both *Roseibium* (*Labrenzia)* ASVs (ASV6 and ASV27), and the *Devosia* ASV (ASV18) all increased in relative abundance with the addition of 4-hydroxybenzoic acid with and without algae present (Figure 6). The fold change of the increase for these ASVs was greater in the algal-free controls compared to algal presence and algal spent medium. Notably the ASV6, ASV18, and ASV23 sequences were exact matches to the 16S-V4 regions of the bacterial isolates *Roseibium* 13C1, *Thalassospira* 13M1, and *Devosia* B7WZ used in the experiments above. The *Thalassospira* ASVs (ASV7 and ASV23) were the most responsive to this metabolite, with bias corrected log2 fold changes of 2.6 and 2.1 respectively in the algae-free condition, and 1.5 and 0.4 respectively with the algae present. Three ASVs (ASV5, ASV10, and ASV15) had divergent responses to 4-hydroxybenzoic acid, increasing under one algal condition and decreasing under another. Three other ASVs (ASV 25, ASV28, and ASV29) decreased in relative abundance with 4-hydroxybenzoic acid addition with algae present and in algal-free controls, while ASV35 decreased in abundance with either algae present or spent medium, but not in controls. Other ASVs either did not change significantly in relative abundance in response to 4-hydroxybenzoic acid addition, or their relative abundance changed only under one algal condition.

**Figure 6.**
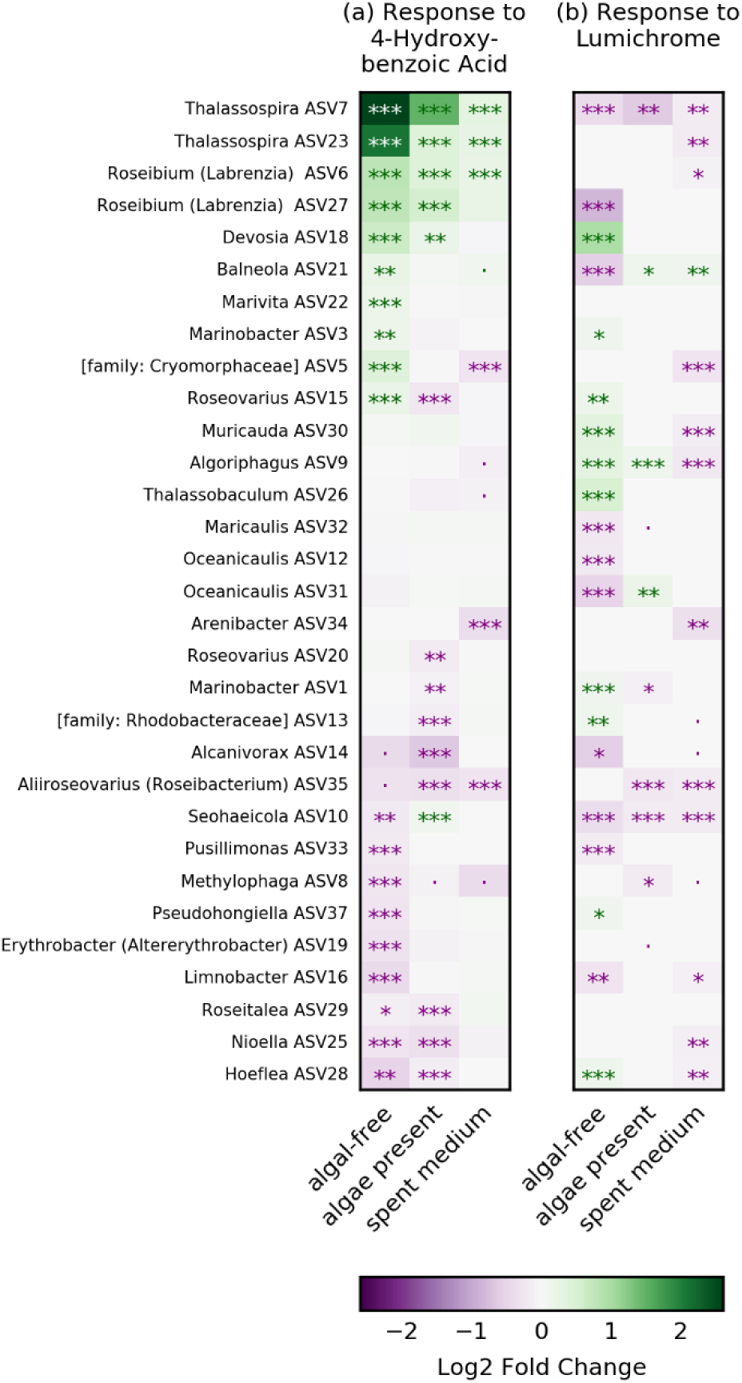
Response of individual ASVs to the addition of exogenous metabolites compared to no metabolite addition. (a) Response to 4-hydroxybenzoic acid. (b) Response to lumichrome. Rows represent ASVs. Columns represent base growth condition (algal-free control, algae present, or algal spent medium). ASV order is the same in (a) and (b) to allow comparison and determined by response to 4-hydroxybenzoic acid. Heatmaps show increases (green) and decreases (purple) in relative abundance of each ASV (rows) with the addition of metabolites under each of the algal conditions (columns). Statistically significant changes in relative abundance are indicated based on the adjusted p-values with the significance codes: 0 ‘***’ 0.001 ‘**’ 0.01 ‘*’ 0.05 ‘.’ 0.1 ‘ ’ 1. For individual comparisons for each ASV under each condition with each metabolite, refer to Supplementary Figures S7, and S8.

Lumichrome addition also impacted relative abundances of ASVs, but the responses differed from those to 4-hydroxybenzoic acid (Figure 6). The magnitude of the strongest responses to lumichrome were not as large as those observed with 4-hydroxybenzoic acid). The maximum bias corrected log2 fold change observed for lumichrome was 0.96, in comparison to 2.6 for 4-hydroxybenzoic acid. No ASVs consistently increased with lumichrome addition across algal treatments. However, two ASVs, *Thalassospira* ASV7 and *Seohaeicola* ASV10, decreased in relative abundance with the addition of lumichrome under all three algal conditions.

## DISCUSSION

In this study we investigated how *P. tricornutum* exometabolites influence an ecologically relevant bacterial community. By identifying and characterizing the impacts of specific metabolites on both representative bacterial isolates and on microbial communities with and without algae or spent medium, we can begin to tease apart the roles of metabolites, algal presence, and bacterial interactions in these systems. Our temporal analysis illustrates how the algal exometabolome has the potential to modulate bacterial ecophysiology as a function of algal growth.

Our findings support the hypothesis that algal exudates can be an important driver of microbial community composition in algal-bacterial systems, but also indicate the importance of other factors that arise with algal presence. We found that algal exudates (*P. tricornutum* spent medium from late-log) shifted the microbial community toward the community composition with algae present, both at the overall community level and at the individual ASV level. However, the microbial communities with algal spent medium still differed from those with algae present, and some individual ASVs had divergent responses to algal spent medium and algal presence. Several important factors may contribute to these differences. First, the presence of the bacterial community can alter algal physiology, which can be expected to result in shifts in algal exudation. Introduction of a bacterial community has been shown to affect algal gene expression in comparison to axenic controls, indicating that algae are actively responding to and potentially modulating the microbial community (14). Other factors arising from algal presence, such as bacterial attachment to algal cells (18), exchange of gases and volatiles (33), the phycosphere microenvironment around algal cells (4), and temporal dynamics (10) likely also contribute to differences between the microbial communities grown with spent medium and those grown with algae. Previous work has shown that attachment can affect both algal carbon fixation and bacterial uptake of algal carbon (18). Thus, the composition of algal exometabolites experienced by bacteria growing with algae likely differs from that of the spent medium from axenic algae. Previous work in our lab investigated bacterial responses to *P. tricornutum* exudates using a porous microplate to physically separate algae and bacteria but allow metabolite exchange (34). Using one of the same bacterial enrichment cultures used here, that study identified four bacterial genera (*Marinobacter*, *Oceanicaulis*, *Algoriphagus*, and *Muricauda*) that responded positively to *P. tricornutum* metabolites diffusing through the hydrogel. In the present study, ASVs from those same genera responded positively to algal spent medium (Figure 2). Thus, both metabolite composition and spatial organization of algae and bacteria influence relative abundances within a microbiome.

Comparing exometabolome compositions over time revealed variation in accumulation patterns of different metabolites over algal growth, which may indicate one factor influencing bacterial populations. Algal-associated microbial communities have been shown to vary with algal growth, for instance across different stages of an algal bloom (35). Previous work investigating algal-bacterial interactions across the diel cycle also found changes in metabolome composition and transcriptomes from one day cycle to the next (10), indicating the importance of the longer time scales that we address here. Both the endometabolome and exometabolome of the marine cyanobacterium *Synechococcus elongatus* also vary over days (36). Further, the extent of organic nitrogen transfer from *P. tricornutum* to bacteria has been shown to vary between bacterial taxa in co-culture studies (19), and thus temporal changes in exudation of nitrogen containing metabolites, which include the majority of metabolites identified here, may influence microbial community dynamics.

Our study shows that an individual metabolite can predictably shift bacterial community composition in the context of both a complex exometabolome (algal spent medium), and in the presence of an algal host. For two isolates (*Thalassospira 13M1* and *Roseibium 13C*), 4-hydroxybenzoic acid acted as a primary organic carbon source, and the corresponding ASVs responded positively to both its addition and, except for one ASV (*Roseibium(Labrenzia))*, to algal spent medium. 4-hydroxybenzoic acid is an organic acid exuded throughout *P. tricornutum* growth, and is also released by *Synechococcus elongatus* (36), suggesting that it may be commonly found in algal exometabolomes. Our previous work quantifying assimilation of algal carbon and nitrogen classified both of these isolates as metabolizing only algal carbon and suggesting specialization for small algal-derived carbon compounds (19). Our results support this classification and demonstrate one such metabolite which gives them a selective advantage when provided to a *P. tricornutum* microbial community. Another study found that compositions of microbial communities grown on individual metabolites were somewhat predictive of compositions of communities grown on suites of five metabolites combined (16), which supports the role we found for algal metabolites in encouraging particular taxa. Genome content also supported our identification of these specialized algal carbon users. We found that the three genera of ASVs with strong positive responses to 4-hydroxybenzoic acid (*Thalasosspira*, *Rosibium* (*Labrenzia*), and *Devosia*), corresponded to the only three of our bacterial isolates with matched ASV’s whose genomes contained the *pobA* gene (Supplementary Table S7). The *pobA* gene encodes for 4-hydroxybenzoate 3-monooxygenase, which converts 4-hydroxybenzoate to protocatechuate, an important entry to the TCA cycle (38). This suggests that *pobA* may convey a competitive advantage within a complex community when 4-hydroxybenzoic acid is readily available.

While 4-hydroxybenzoic acid exemplifies selective growth promotion as a potential mechanism of algal impacts on microbial communities, inhibition effects, while not as straightforward, are also an important factor in modulating microbial communities. Taxa previously shown to specialize in algal carbon assimilation (19) generally responded positively to algal spent medium, but negatively to algal presence, and in some cases to lumichrome addition. Along with *Thalassospira* and *Roseibium (Labrenzia)* this included *Marinobacter*, which did not grow on 4-hydroxybenzoic acid, nor respond positively to its addition, but did respond negatively to algal presence but not algal spent medium. The is notable since most other ASVs responded in the same direction to both algal presence and spent medium. This response suggests that other factors associated specifically with algal presence selectively reduce the growth of these taxa in a complex algal-bacterial system, and that excess levels of an algal exometabolite to which they are specialized (4-hydroxybenzoic acid) can overcome this inhibition. Competition for inorganic nutrients may contribute to this effect. A previous study using porous microplates with gradients of inorganic nutrients and algal DOC found that growth of *Marinobacter* (same isolate used here) was dependent on access to inorganic nutrients along with algal DOC (34). However, that study also found that when a bacterial enrichment community (one of the same enrichments used here) was grown in adjacent porous microplate wells to *P. tricornutum*, its response was similar to this study’s spent medium response (increases in *Marinobacter*, *Oceanicaulus* and *Algoriphagus*, and a decrease in *Alcanivorax* relative abundance), not the algal presence response (40).This suggests that direct physical interaction is required for the inhibition effect observed in this study (34). Other work has shown that antimicrobial fatty acids from *P. tricornutum* cell pellets selectively inhibited some bacterial taxa (42), suggesting one potential mechanism of selective controls of bacterial growth tied to algal presence. Alternatively, direct interactions between the algae and other bacterial taxa could in turn impact interactions between those taxa and the algal carbon specialists.

The effects of lumichrome addition illustrate response to a different class of metabolite that may not be directly involved with nutrition and highlight interactions that may involve other factors such as signaling and microbe-microbe interactions. Lumichrome, a vitamin derivative, did have significant effects on microbial abundances. However, it was not an effective growth substrate for the bacterial isolates, and, perhaps because of this, our previous characterization of the isolates’ exudate carbon and nitrogen usage (19) was not predictive of response. Lumichrome has previously been shown to stimulate the growth of *P. tricornutum* and of other microalgae (11, 44), and to change the endometabolite composition of *Chlorella sorokiniana* (44). This suggests that differences in microbial abundances occurring when algae are present may be indirect effects due to lumichrome altering algal physiology which in turn affects the bacteria. Lumichrome is a degradation product of riboflavin (vitamin B2) and can also act as a bacterial quorum sensing mimic (45). In either of these roles, lumichrome may also have a more direct effect on the growth and physiology of bacterial community members.

Our results demonstrate the multifaceted role of algal extracellular metabolites in shaping algal-associated bacterial communities. Using a relevant suite of bacterial isolates and an enrichment community containing similar taxa, we identified specific algal metabolites (4-hydroxybenzoic acid and lumichrome) that influenced individual taxa and showed that a selective bacterial growth substrate (4-hydroxybenzoic acid) represents one mechanism by which algal exudates can modulate the microbial community. Our work also provides evidence for the presence of additional controls requiring algal presence that selectively keep taxa specializing in algal carbon exometabolite consumption in check relative to taxa with other ecological strategies. We also show the dynamics of exometabolite production, with important implications for different bacterial with different carbon and nitrogen utilization strategies and substrate preferences. This work advances our understanding of the algal exometabolome and its importance in algal-bacterial interactions and microbial community composition.

## ACKNOWLEDGEMENTS

This research was supported by the LLNL microBiospheres Scientific Focus Area, funded by the U.S. Department of Energy Office of Science, Office of Biological and Environmental Research Genomic Science program under FWP SCW1039. This work was performed under the auspices of the U.S. Department of Energy by Lawrence Livermore National Laboratory under Contract DE-AC52-07NA27344. LLNL IM release number LLNL-JRNL-836013-DRAFT.

## COMPETING INTERESTS

The authors declare no competing financial interests.

## DATA AVAILABILITY

Sequencing data are deposited in the NCBI Sequence Read Archive under BioProject number PRJNA803592. Metabolomics LC-MS/MS raw data and metadata are deposited in the NIH Common Fund’s National Metabolomics Data Repository (supported by NIH grant U2C-DK119886) website, the Metabolomics Workbench, https://www.metabolomicsworkbench.org under Project ID PR001317 and Study ID ST002077, and can be accessed directly via the Project DOI: http://dx.doi.org/10.21228/M8GH6P.

## Notes

https://www.ncbi.nlm.nih.gov/sra/PRJNA803592

http://dx.doi.org/10.21228/M8GH6P

### Competing Interest Statement

The authors have declared no competing interest.

## REFERENCES

1. Falkowski P. OCEAN SCIENCE The power of plankton. Nature. 2012;483(7387):S17–S20.

2. Chisti Y. Biodiesel from microalgae. Biotechnology Advances. 2007;25(3):294–306.

3. Mata TM, Martins AA, Caetano NS. Microalgae for biodiesel production and other applications: A review. Renewable & Sustainable Energy Reviews. 2010;14(1):217–32.

4. Seymour JR, Amin SA, Raina JB, Stocker R. Zooming in on the phycosphere: the ecological interface for phytoplankton-bacteria relationships. Nature Microbiology. 2017;2(7):12.

5. Yao S, Lyu S, An Y, Lu J, Gjermansen C, Schramm A. Microalgae-bacteria symbiosis in microalgal growth and biofuel production: a review. Journal of Applied Microbiology. 2019;126(2):359–68.

6. Lian J, Wijffels RH, Smidt H, Sipkema D. The effect of the algal microbiome on industrial production of microalgae. Microbial Biotechnology. 2018;11(5):806–18.

7. Seyedsayamdost MR, Case RJ, Kolter R, Clardy J. The Jekyll-and-Hyde chemistry of Phaeobacter gallaeciensis. Nature Chemistry. 2011;3(4):331–5.

8. Wang H, Tomasch J, Jarek M, Wagner-Dobler I. A dual-species co-cultivation system to study the interactions between Roseobacters and dinoflagellates. Frontiers in Microbiology. 2014;5.

9. Thornton DCO. Dissolved organic matter (DOM) release by phytoplankton in the contemporary and future ocean. European Journal of Phycology. 2014;49(1):20–46.

10. Uchimiya M, Schroer W, Olofsson M, Edison AS, Moran MA. Diel investments in metabolite production and consumption in a model microbial system. The ISME Journal. 2021.

11. Brisson V, Mayali X, Bowen B, Golini A, Thelen M, Stuart RK, et al. Identification of Effector Metabolites Using Exometabolite Profiling of Diverse Microalgae. mSystems. 2021;6(6):e00835–21.

12. Ferrer-Gonzalez FX, Widner B, Holderman NR, Glushka J, Edison AS, Kujawinski EB, et al. Resource partitioning of phytoplankton metabolites that support bacterial heterotrophy. Isme Journal. 2021;15(3):762–73.

13. Becker JW, Berube PM, Follett CL, Waterbury JB, Chisholm SW, DeLong EF, et al. Closely related phytoplankton species produce similar suites of dissolved organic matter. Frontiers in Microbiology. 2014;5:14.

14. Shibl AA, Isaac A, Ochsenkuhn MA, Cardenas A, Fei C, Behringer G, et al. Diatom modulation of select bacteria through use of two unique secondary metabolites. Proceedings of the National Academy of Sciences of the United States of America. 2020;117(44):27445–55.

15. Kimbrel JA, Samo TJ, Ward C, Nilson D, Thelen MP, Siccardi A, et al. Host selection and stochastic effects influence bacterial community assembly on the microalgal phycosphere. Algal Research-Biomass Biofuels and Bioproducts. 2019;40:10.

16. Fu H, Uchimiya M, Gore J, Moran MA. Ecological drivers of bacterial community assembly in synthetic phycospheres. Proceedings of the National Academy of Sciences of the United States of America. 2020;117(7):3656–62.

17. Seyedsayamdost MR, Wang RR, Kolter R, Clardy J. Hybrid Biosynthesis of Roseobacticides from Algal and Bacterial Precursor Molecules. Journal of the American Chemical Society. 2014;136(43):15150–3.

18. Samo TJ, Kimbrel JA, Nilson DJ, Pett-Ridge J, Weber PK, Mayali X. Attachment between heterotrophic bacteria and microalgae influences symbiotic microscale interactions. Environmental Microbiology. 2018;20(12):4385–400.

19. Mayali X, Samo T, Kimbrel J, Stuart RK, Morris M, Rolison K, et al. Diatom-associated bacteria exhibit contrasting ecological strategies for growth and metabolite exchange. In prep. 2022.

20. Chorazyczewski AM, Huang IS, Abdulla H, Mayali X, Zimba PV. The Influence of Bacteria on the Growth, Lipid Production, and Extracellular Metabolite Accumulation by Phaeodactylum tricornutum (Bacillariophyceae). Journal of Phycology. 2021;57(3):931–40.

21. Berges JA, Franklin DJ, Harrison PJ. Evolution of an artificial seawater medium: Improvements in enriched seawater, artificial water over the last two decades. Journal of Phycology. 2001;37(6):1138–45.

22. Harrison PJ, Waters RE, Taylor FJR. A BROAD-SPECTRUM ARTIFICIAL SEAWATER MEDIUM FOR COASTAL AND OPEN OCEAN PHYTOPLANKTON. Journal of Phycology. 1980;16(1):28–35.

23. Guillard RR, Ryther JH. STUDIES OF MARINE PLANKTONIC DIATOMS.1. CYCLOTELLA NANA HUSTEDT, AND DETONULA CONFERVACEA (CLEVE) GRAN. Canadian Journal of Microbiology. 1962;8(2):229-&.

24. Zhalnina K, Louie KB, Hao Z, Mansoori N, da Rocha UN, Shi SJ, et al. Dynamic root exudate chemistry and microbial substrate preferences drive patterns in rhizosphere microbial community assembly. Nature Microbiology. 2018;3(4):470–80.

25. Swift CL, Louie KB, Bowen BP, Olson HM, Purvine SO, Salamov A, et al. Anaerobic gut fungi are an untapped reservoir of natural products. Proceedings of the National Academy of Sciences of the United States of America. 2021;118(18).

26. Bowen BP, Northen TR. Dealing with the Unknown: Metabolomics and Metabolite Atlases. Journal of the American Society for Mass Spectrometry. 2010;21(9):1471–6.

27. Yao YS, Sun T, Wang T, Ruebel O, Northen T, Bowen BP. Analysis of Metabolomics Datasets with High-Performance Computing and Metabolite Atlases. Metabolites. 2015;5(3):431–42.

28. Parada AE, Needham DM, Fuhrman JA. Every base matters: assessing small subunit rRNA primers for marine microbiomes with mock communities, time series and global field samples. Environmental Microbiology. 2016;18(5):1403–14.

29. Apprill A, McNally S, Parsons R, Weber L. Minor revision to V4 region SSU rRNA 806R gene primer greatly increases detection of SAR11 bacterioplankton. Aquatic Microbial Ecology. 2015;75(2):129–37.

30. Callahan BJ, McMurdie PJ, Rosen MJ, Han AW, Johnson AJA, Holmes SP. DADA2: High-resolution sample inference from Illumina amplicon data. Nature Methods. 2016;13(7):581-+.

31. McMurdie PJ, Holmes S. phyloseq: An R Package for Reproducible Interactive Analysis and Graphics of Microbiome Census Data. Plos One. 2013;8(4):11.

32. Lin H, Das Peddada S. Analysis of compositions of microbiomes with bias correction. Nature Communications. 2020;11(1):11.

33. Zuo ZJ. Why Algae Release Volatile Organic Compounds - The Emission and Roles. Frontiers in Microbiology. 2019;10:7.

34. Kim H, Kimbrel JA, Vaiana CA, Wollard JR, Mayali X, Buie CR. Bacterial response to spatial gradients of algal-derived nutrients in a porous microplate. Isme Journal. 2021:10.

35. Zhou J, Chen GF, Ying KZ, Jin H, Song JT, Cai ZH. Phycosphere Microbial Succession Patterns and Assembly Mechanisms in a Marine Dinoflagellate Bloom. Applied and Environmental Microbiology. 2019;85(15):17.

36. Fiore C, Longnecker K, Soule M, Kujawinski E. Release of ecologically relevant metabolites by the cyanobacterium Synechococcus elongatusCCMP 1631. Environmental Microbiology. 2015;17(10):3949–63.

37. . !!! INVALID CITATION !!! (34).

38. Harwood CS, Parales RE. The beta-ketoadipate pathway and the biology of self-identity. Annual Review of Microbiology. 1996;50:553–90.

39. Cho JY, Moon JH, Seong KY, Park KH. Antimicrobial activity of 4-hydroxybenzoic acid and trans 4-hydroxycinnamic acid isolated and identified from rice hull. Bioscience Biotechnology and Biochemistry. 1998;62(11):2273–6.

40. Arcand MM, Schneider KD. Plant- and microbial-based mechanisms to improve the agronomic effectiveness of phosphate rock: a review. Anais Da Academia Brasileira De Ciencias. 2006;78(4):791–807.

41. . !!! INVALID CITATION !!! (36).

42. Desbois AP, Mearns-Spragg A, Smith VJ. A Fatty Acid from the Diatom Phaeodactylum tricornutum is Antibacterial Against Diverse Bacteria Including Multi-resistant Staphylococcus aureus (MRSA). Marine Biotechnology. 2009;11(1):45–52.

43. . !!! INVALID CITATION !!! (35).

44. Lopez BR, Palacios OA, Bashan Y, Hernandez-Sandoval FE, de-Bashan LE. Riboflavin and lumichrome exuded by the bacterium Azospirillum brasilense promote growth and changes in \metabolites in Chlorella sorokiniana under autotrophic conditions. Algal Research-Biomass Biofuels and Bioproducts. 2019;44.

45. Rajamani S, Bauer WD, Robinson JB, Farrow JM, Pesci EC, Teplitski M, et al. The Vitamin Riboflavin and Its Derivative Lumichrome Activate the LasR Bacterial Quorum-Sensing Receptor. Molecular Plant-Microbe Interactions. 2008;21(9):1184–92.

