## Supplementary Information for "Dynamic *Phaeodactylum tricornutum* Exometabolites Shape Surrounding Bacterial Communities"

### SUPPLEMENTARY METHODS

#### Organisms

We used the model diatom *P. tricornutum* strain CCMP 2561. The bacterial isolates used in this study were isolated in our lab and have been described elsewhere (1, 2). For this study we used ten phycosphere bacterial isolates from *P. tricornutum* phycosphere enrichment cultures and two non-phycosphere bacterial isolates from sea water. Information on the bacterial isolates origins and identifications is summarized in Table 1. The enrichment culture used in this study was from the same set of phycosphere enrichments used to obtain the bacterial isolates and has been described elsewhere (3, 4).

#### Growth medium

Experiments were conducted using Enriched Saltwater Artificial Water (ESAW) with F/2 medium trace metals and vitamins, which has been described previously (5-8). The medium contained salts (21.194 g/L NaCl, 3.55 g/L $Na_2SO_4$, 0.599 g/L KCl, 0.174 g/L $NaHCO_3$, 0.0863 g/l KBr, 0.023 g/L $H_3PO_4$, 0.0028 g/L NaF, 9.592 g/L $MgCl_2\cdot6H_2O$, 1.344 g/L $CaCl_2\cdot2H_2O$, 0.0218 g/L $SrCl_2\cdot6H_2O$, and 0.03 g/L $Na_2SiO_3\cdot9H_2O$), phosphate (0.005 g/L $NaH_2PO_4.H_2O$), trace metals (0.023 mg/L $ZnSO_4\cdot7H_2O$, 0.152 mg/L $MnSO_4\cdot H_2O$, 0.0073 mg/L $NaMoO_4\cdot2H_2O$, 0.014 mg/L $CoSO_4\cdot7H_2O$, 0.0068 mg/L $CuCl_2\cdot2H_2O$, 4.6 mg/L $Fe(NH_4)_2(SO_4)_2\cdot6H_2O$, and 4.4 mg/L $Na_2EDTA\cdot2H_2O$), and vitamins (0.0135 mg/L cyanocobalamin, 0.0025 mg/L biotin, and 0.0335 mg/L thiamine). For bacterial isolate growth experiments, nitrogen was provided as equimolar concentrations of nitrate and ammonium (0.0375 g/L $NaNO_3$ and 0.0236 g/L $NH_4Cl$). For all other experiments, nitrogen was provided as nitrate (0.075 g/L $NaNO_3$).

**Algal exometabolomics experiment**

***Algal growth.*** Axenic algal cultures were grown 125 mL glass Erlenmeyer flasks in 50 mL ESAW medium. Prior to the experiment, glass flasks were baked at 500 °C for two hours to remove residual organic matter. Flasks were incubated with shaking at 90 rpm and 22 °C with a 12-hour day/night cycle and a daytime illumination of 3500 lux. Twenty replicate flasks were inoculated at the start of the experiment, each with two mL from a one-week-old culture of *P. tricornutum*. Algal growth was monitored by measuring chlorophyll A fluorescence of 1 mL subsamples using a Trilogy Fluorometer [Turner Designs, San Jose, CA, USA] with the Chlorophyll A In-Vivo Module. Sampling times are indicated on the growth curve in Supplementary Figure S1.

***Sample collection.*** At each metabolite collection time point (0, 3, 6, 9, and 12 days), four flasks were destructively sampled for metabolomics analysis of spent medium and cell pellets as previously described (8). Briefly, cell pellets were collected by centrifugation at 5000 g and 4 °C for 8 minutes. The supernatant was filtered through a 0.45 µm pore size filter and frozen at -80 °C. Cell pellets were resuspended in 1 mL 0.2 µm filtered sterile water, frozen at -80 °C, and lyophilized. Lyophilized cells were disrupted by vortexing with a steel ball three times for 5 seconds each, resuspended in 50 mL 0.2 um filtered sterile water, filtered through a 0.45 µm pore size filter and frozen at -80 °C.

***Metabolite extraction.*** For solid phase extraction of metabolites from frozen samples (filtered spend medium and filtered water extracts of disrupted cell pellets), samples were thawed and 30 mL of each sample was acidified by adding 300 µL of 1 M HCl. Bond Elut PPL columns

[Agilent, Santa Clara, CA, USA] were prepared by rinsing with 3 mL HPLC grade methanol followed by with 6 mL ultrapure water according to the manufacturer's instructions. Acidified samples were passed through the prepared columns. Columns were rinsed with 3 mL of 0.01 M HCl and allowed to air dry of 5 minutes. Metabolites were eluted with 1 mL HPLC grade methanol. Methanol extracts were dried in a vacuum centrifuge, and dried extracts were resuspended and prepared for analysis as previously described (8-10). Briefly, extracts were resuspended in 10 µL methanol containing $^{13}C$ and $^{15}N$ labeled matrix control standards, filtered through a 0.2 um pore size filter, and transferred to an autosampler vial for metabolomic analysis.

***Metaoblomics analysis.*** Metabolomes were analyzed with liquid chromatography tandem mass spectrometry (LC-MS/MS). LC-MS/MS instrumentation and parameters are detailed in Supplementary Table S1. Metabolite identifications were obtained using Metabolite Atlas (https://github.com/biorack/metatlas) to identify features with *m/z*, retention time, and fragmentation spectra matched to a library of analytical standards run in our lab under the same LC-MS/MS analysis conditions (11, 12).

**Bacterial isolate growth assays**

Twelve selected exometabolites were tested for their ability to support growth as the carbon source for each of each of twelve bacterial isolates. Bacterial isolate growth assays were conducted in flat bottomed 96 well plates. For each metabolite tested, ESAW growth medium was prepared with the added metabolite at a total carbon concentration of 60 ppm. This concentration was chosen as approximately ten times the previously observed TOC concentration in *P. tricornutum* spent medium (8), representing total DOC, in order to stimulate sufficient growth for detection. Each metabolite was tested with each bacterial isolate in triplicate wells with replicate positions randomized across plates. Negative controls were also included, with no metabolites added to the medium. Plates were incubated at room temperature (24 °C) and shaken at 90 rpm. Bacterial growth was measured as OD 600 in a Cytation5 plate reader [BioTek, Winooski, VT, USA]. Growth was measured daily for eight days.

**Microbial community response experiments**

***Growth conditions.*** Three algal conditions were tested: algae present, algal spent medium, and algal-free control. Experiments were conducted in 125 mL flasks prepared as described above for the metabolomics experiment. For the algae present condition, flasks were prepared with ESAW medium and inoculated with 2 mL of one-week-old *P. tricornutum* culture at the same time as the bacterial inoculation. For the algal spent medium condition, flasks were prepared with spent medium from a one-week-old *P. tricornutum* culture that was filtered through a 0.2 micron filter to remove algal cells. Phosphate and nitrate were replenished in the algal spent medium by adding 0.005 g/L $NaH_2PO_4.H_2O$ and 0.075 g/L $NaNO_3$ from 1000x stock solutions prior to inoculation with bacteria. Algal-free control flasks were prepared with ESAW medium without any addition of alga, algal spent medium, or other carbon source beyond what was present in the base medium.

***Metabolite additions.*** For the added metabolite experiments, the same three algal conditions as above were prepared with the addition of 0.01 mM of the metabolite to be tested. This concentration, lower than that used in the bacterial isolate experiment above, was selected for two reasons: (A) so that the added metabolite represented a small change and did not overwhelm the algal metabolites present in the algal spent medium and algae present conditions, and (B) because previous work with lumichrome indicated that this concentration had effects on *P. tricornutum* growth (8).

***Inoculation.*** To obtain bacterial inoculum for this experiment, two previously described phycosphere enrichment cultures were combined (3, 4). One enrichment was maintained with *P. tricornutum,* while the other was maintained with *Microchloropsis salina*. Enrichments were grown in ESAW medium for one week. Cultures were then passed through a 45 mm diameter, 0.8 um pore size Supor membrane filter [Pall Corporation, Port Washington, NY, USA], to remove algal cells but allow bacterial cells to pass through. This filtrate was used to inoculate all flasks with 2 mL (1 mL from each enrichment). Once inoculated, flasks were incubated as described above for metabolomics experiments.

***Sample collection and DNA extraction.*** Samples were collected after 6, 8, and 10 days of incubation. Twenty-five mL was collected from each flask, and flasks were replenished with 25 mL of the appropriate fresh medium for that flask. From the 25 mL sample, one mL was fixed for flow cytometry measurements by adding 100 µL 37% formaldehyde. The remaining sample was filtered onto a 45 mm diameter, 0.2 µm pore size Supor membrane filter [Pall Corporation, Port Washington, NY, USA], and filters were frozen at -80 °C until DNA extraction. DNA extractions were conducted using the NucleoSpin 96 Tissue Kit [Macherey-Nagel, Düren, Germany] with modifications to the vacuum processing protocol. Filters were divided in half, and each half was processed separately for the first steps of extraction. Half filters were pre-lysed with 345 µL T1 buffer and 15 µL lysozyme (50 mg/mL) at 37 °C for 30 minutes. Proteinase K (50 uL) was added, and samples were incubated at 56 °C in rotating tubes overnight (approximately 16 hours) to mix and lyse. After cell lysis, 400 µL of BQ1 buffer and 400 µL of 100% ethanol were added to each sample, and lysates were transfered to the NucleoSpin Tissue Binding Plate. Halves from the same sample were combined at this point. Samples were vaccumed through the manifold at -0.2 bar for 5 minutes. Membranes were washed three times (first wash: 600 µL of BW buffer, second wash: 900 µL of B5 buffer, third wash: 900 µL of B5 buffer) and vacuumed at -0.2 bar for 5 minutes after each wash. Membranes were dried by vacuuming at -0.6 bar for 10 minutes. Samples were eluted 2x into microtubes by adding 60 µL of BE buffer (preheated to 70 °C), incubating at room temperature for 3 to 5 minutes, and vacuuming at -0.4 bar for 2 minutes. DNA concentrations were quantified using the Invitrogen Qubit dsDNA HS assay kit [Thermo Fischer Scientific, Waltham, MA, USA] and stored at -20 °C until sequencing.

***Amplicon sequencing and analysis.*** Sequencing was performed through Laragen, Inc. [Culver City, CA, USA], using the earth microbiome project protocol. The 515F (GTGYCAGCMGCCGCGGTAA) (13) and 806R (GGACTACNVGGGTWTCTAAT) (14) primers were used to amplify and sequence the 16S-V4 region. Sequencing was performed on an Illumina MiSeq [Illumina, San Diego, CA, USA] using the MiSeq V2 PE150 reagent kit. Adapter sequences were trimmed from sequencing reads using cutadapt, and trimmed reads were processed using the DADA2 package (version 1.20.0) in R (version 4.1.2) to assess read quality, filter reads, merge paired reads and remove chimeras, and generate an amplicon sequence variant (ASV) table (15). ASV taxonomy was assigned using both the RDP database (version 18) and the Silva database (version 138). Where identifications conflicted, the RDP identification is given with the Silva identification in parentheses. Microbial community analysis was conducted using the phyloseq package (version 1.36.0) in R (16). ASVs were filtered to remove chloroplast and mitochondrial sequences, and to remove ASVs that were not detected with at least ten counts in at least four samples. The ANCOM-BC package (version 1.2.2) was used to assess differential abundance and to generate bias corrected ASV counts for further analyses (17).

### REFERENCES FOR SUPPLEMENTARY INFORMATION

1. Mayali X, Samo T, Kimbrel J, Stuart RK, Morris M, Rolison K, et al. Diatom-associated bacteria exhibit contrasting ecological strategies for growth and metabolite exchange. In prep. 2022.

### SUPPLEMENTARY TABLES

**Supplementary Table S1.**

LCMS instrument and parameters for metabolomics analysis. (Excel File)

**Supplementary Table S2.**

PERMANOVA of prokaryotic community composition based on relative abundances of 62 ASVs. Significance codes: 0 ‘***’ 0.001 ‘**’ 0.01 ‘*’ 0.05 ‘.’ 0.1 ‘ ’ 1.

| Sources of Variation | Degrees of Freedom | Sum Of Squares | F statistic | Pr (>F) | Statistical Significance |
|---|---|---|---|---|---|
| algae | 2 | 0.786 | 802.0 | 0.0001 | *** |
| time | 2 | 0.018 | 18.4 | 0.0001 | *** |
| algae : time | 4 | 0.024 | 12.4 | 0.0001 | *** |
| residuals | 36 | 0.018 | | | |
| total | 44 | 0.846 | | | |

**Supplementary Table S3.**

ANOVA of microbial community alpha diversity as measured by the Shannon diversity index. This is a type III ANOVA analyzing algal treatment, time, and their interaction. Significance codes: 0 ‘***’ 0.001 ‘**’ 0.01 ‘*’ 0.05 ‘.’ 0.1 ‘ ’ 1.

| Sources of Variation | Degrees of Freedom | Sum Of Squares | F statistic | Pr (>F) | Statistical Significance |
|---|---|---|---|---|---|
| algae | 2 | 0.704 | 85.1 | < 0.0001 | *** |
| time | 2 | 0.391 | 47.3 | < 0.0001 | *** |
| algae : time | 4 | 0.125 | 7.5 | 0.0002 | *** |

**Supplementary Table S4.**

Metabolite identifications. (Excel file)

**Supplementary Table S5.**

Metabolite quantification. (Excel file)

**Supplementary Table S6.**

PERMANOVA of prokaryotic community composition based on relative abundances of 62 ASVs. Significance codes: 0 ‘***’ 0.001 ‘**’ 0.01 ‘*’ 0.05 ‘.’ 0.1 ‘ ’ 1.

| Sources of Variation | Degrees of Freedom | Sum Of Squares | F statistic | Pr (>F) | Statistical Significance |
|---|---|---|---|---|---|
| algae | 2 | 2.220 | 4766.6 | 0.0001 | *** |
| metabolite | 2 | 0.326 | 699.7 | 0.0001 | *** |
| time | 2 | 0.053 | 112.7 | 0.0001 | *** |
| algae : metabolite | 4 | 0.111 | 119.6 | 0.0001 | *** |
| algae : time | 4 | 0.071 | 76.4 | 0.0001 | *** |
| metabolite : time | 4 | -0.002 | -5.5 | 1 | |
| algae : metabolite : time | 8 | -0.034 | -18 | 1 | |
| residuals | 108 | 0.025 | | | |
| total | 134 | 2.768 | | | |

**Supplementary Table S7.**

Summary of responses of bacterial isolates and congeneric ASVs. (Excel File)

### SUPPLEMENTARY FIGURES

**Supplementary Figure S1**

Algal growth and sampling. Algal growth was monitored as Chlorophyl A fluorescence measured in relative fluorescence units (RFU). Sampling time points for metabolomics are indicated with arrows. Colors indicate different growth phases: yellow = lag phase, green = early log phase, blue = late log phase, purple = stationary phase.

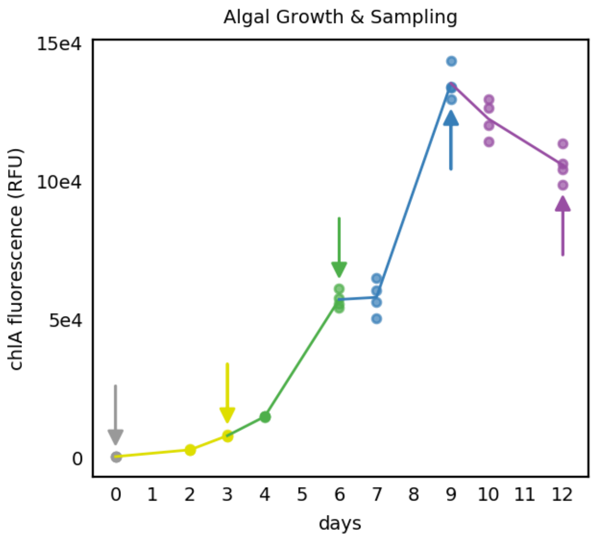

**Supplementary Figure S2.**

Relative abundances of ASVs in individual samples, grouped by conditions and time points. All 13 ASVs with mean relative abundance over 0.1% are shown.

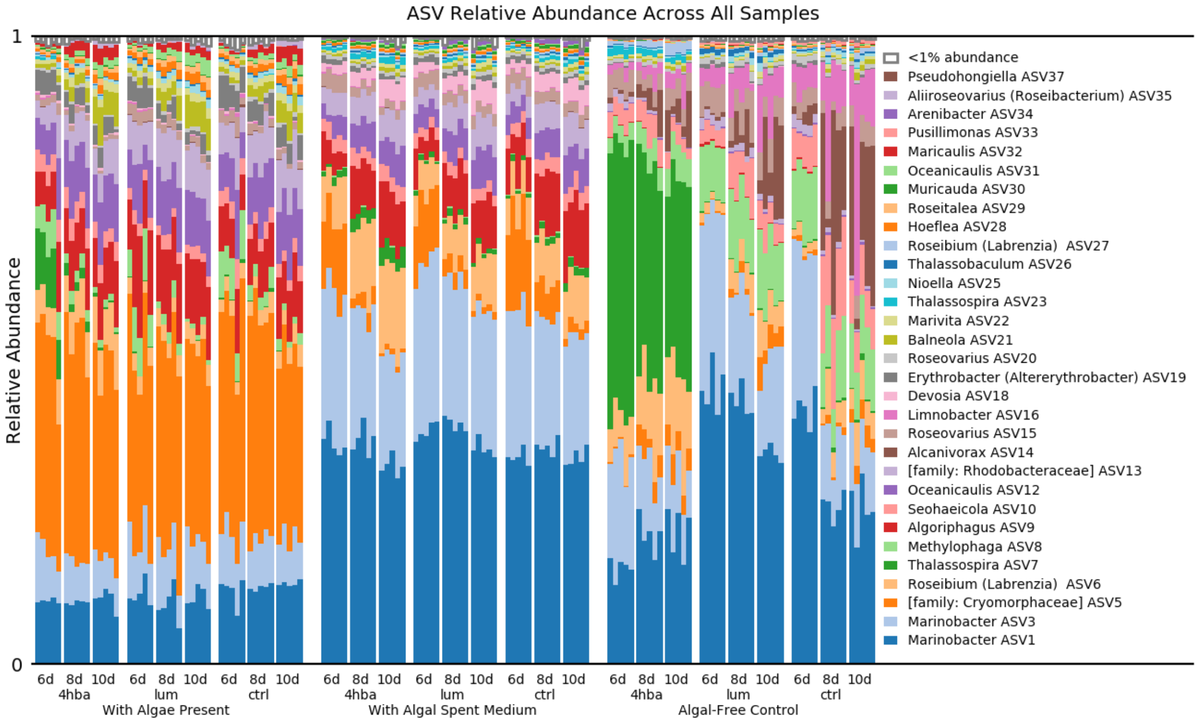

**Supplementary Figure S3.**

Microbial community alpha diversity as measured by Shannon diversity index. (a) Different algal treatments. (b) Trends over time. (c) Algal treatment and time interactions. Letters indicate statistically distinct groups based on the post-hoc Tukey's tests.

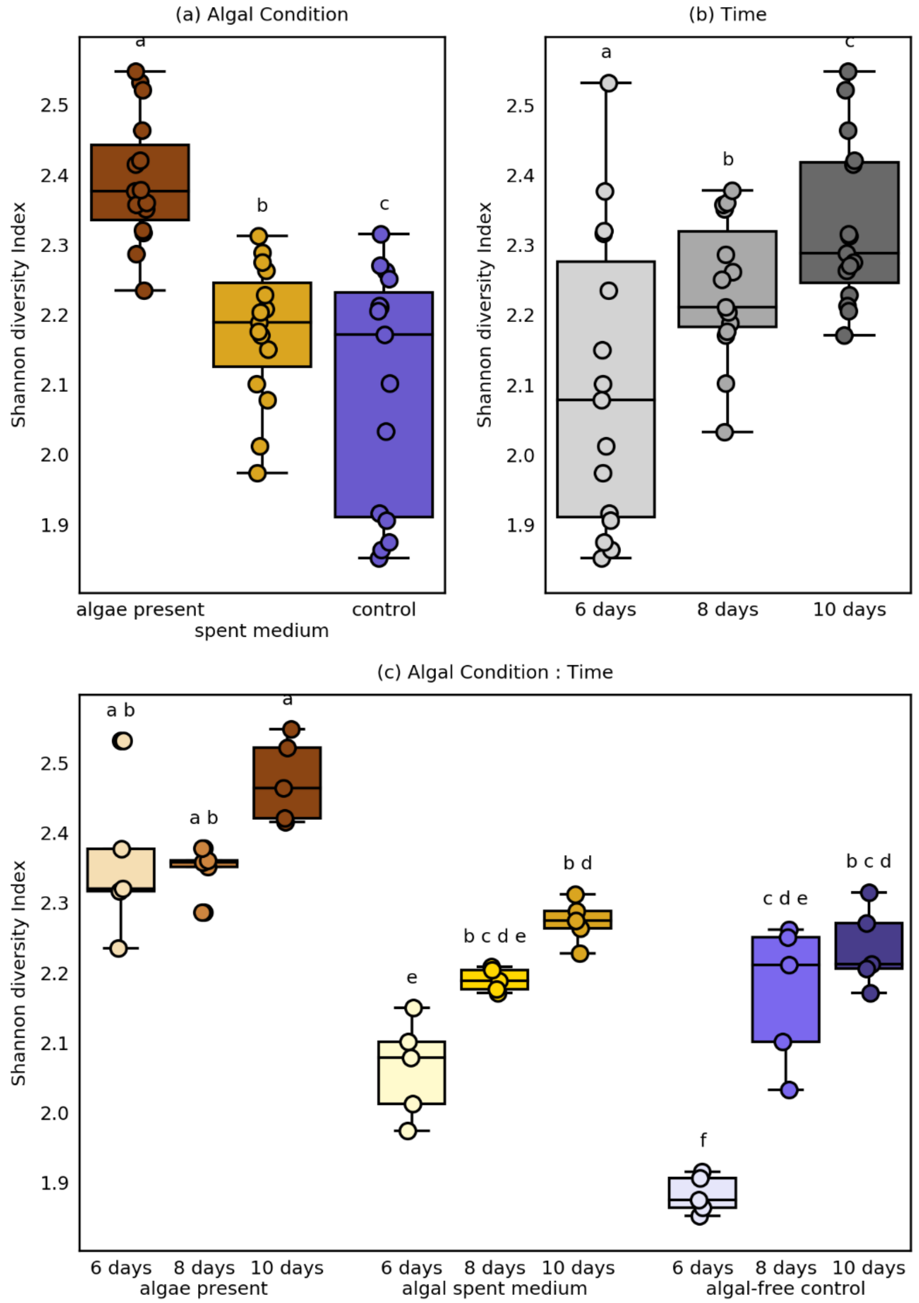

**Supplementary Figure S4.**

Ternary plot showing bacterial ASV responses to algal exudates and algal presence. Each circle represents one ASV, and its position indicates its relative proportion among the three growth conditions (with algae, with algal spent medium, and algal-free control). Circle size is proportional to ASV mean relative abundance. Color of the left half of the circle indicates response to algal spent medium in comparison to algal-free controls. Color of the right half of the circle indicates response to algal presence in comparison to algal-free controls. ASVs above 0.5% mean relative abundance are labeled with ASV IDs.

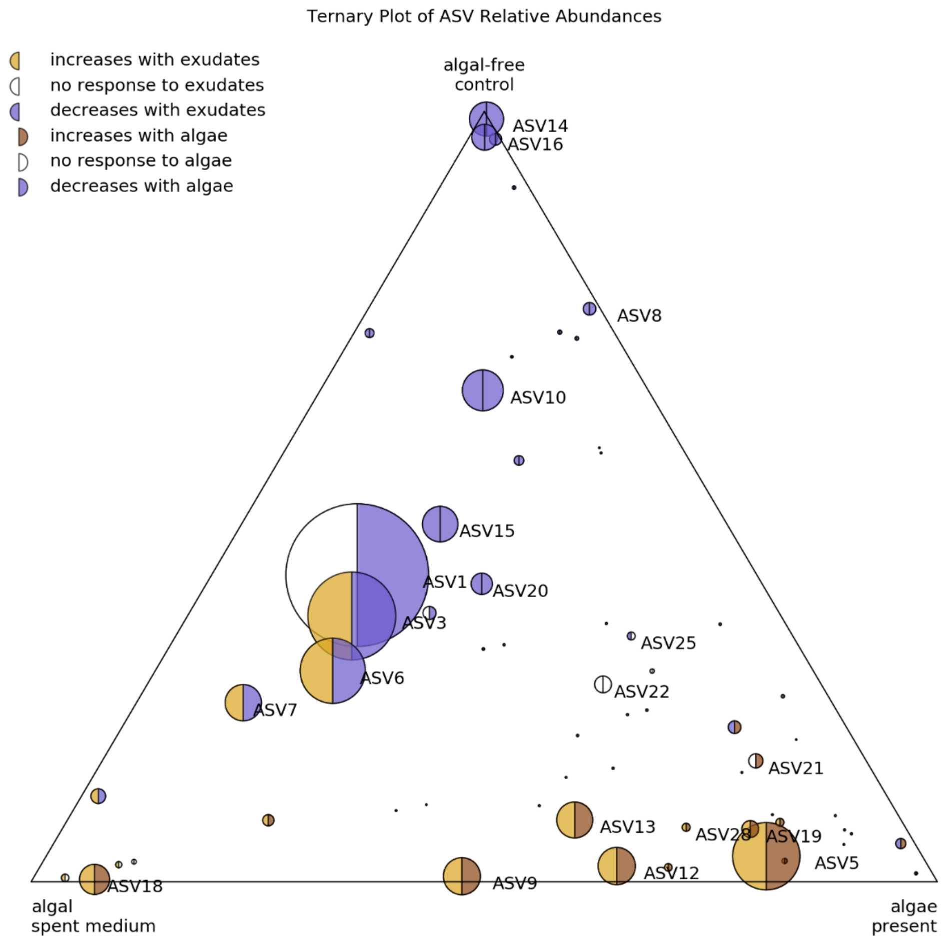

**Supplementary Figure S5.**

Metabolomics heatmap showing 50 identified metabolites' relative intensities across samples. Each column represents one sample, and samples are grouped by sample type and time point as indicated above the plot. Each row represents one metabolite (or indistinguishable isomers). Metabolites are ordered based on hierarchical clustering, as indicated by the dendrogram to the right of the heatmap. Heatmap coloring at each point corresponds to the relative intensity for each metabolite in each sample. Relative intensity is the signal intensity for a given metabolite in a sample divided by the maximum observed signal intensity for that metabolite and is therefore on a scale of 0 to 1.

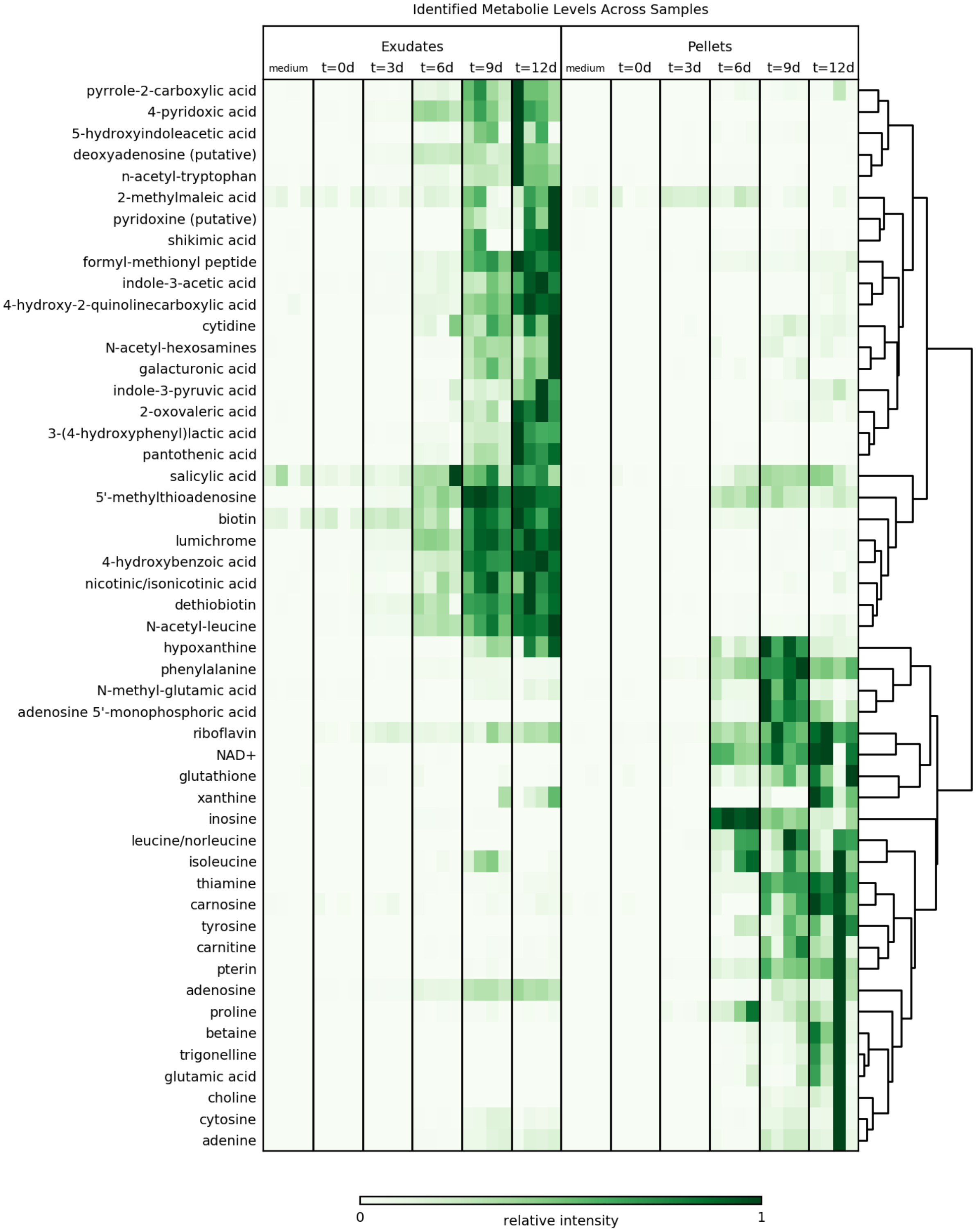
Identified Metabolie Levels Across Samples
Exudates
Pellets
medium
t=0d
t=3d
t=6d
t=9d
t=12d
pyrrole-2-carboxylic acid
4-pyridoxic acid
5-hydroxyindoleacetic acid
deoxyadenosine (putative)
n-acetyl-tryptophan
2-methylmaleic acid
pyridoxine (putative)
shikimic acid
formyl-methionyl peptide
indole-3-acetic acid
4-hydroxy-2-quinolinecarboxylic acid
cytidine
N-acetyl-hexosamines
galacturonic acid
indole-3-pyruvic acid
2-oxovaleric acid
3-(4-hydroxyphenyl)lactic acid
pantothenic acid
salicylic acid
5'-methylthioadenosine
biotin
lumichrome
4-hydroxybenzoic acid
nicotinic/isonicotinic acid
dethiobiotin
N-acetyl-leucine
hypoxanthine
phenylalanine
N-methyl-glutamic acid
adenosine 5'-monophosphoric acid
riboflavin
NAD+
glutathione
xanthine
inosine
leucine/norleucine
isoleucine
thiamine
carnosine
tyrosine
carnitine
pterin
adenosine
proline
betaine
trigonelline
glutamic acid
choline
cytosine
adenine
0
relative intensity
1

249

250

**Supplementary Figure S6.**

Bacterial Isolate growth results for the two metabolites that supported significant growth and for the no metabolite added controls for comparison. Bars indicate the mean optical density for each isolate with that metabolite. Points indicate each of the three individual replicates and are connected by a vertical line indicating the range of the data. Symbols above bars indicate statistically significant increases in optical density compared to the control for that isolate.

Significance codes: 0 '***' 0.001 '**' 0.01 '*' 0.05 '.' 0.1 ' ' 1.

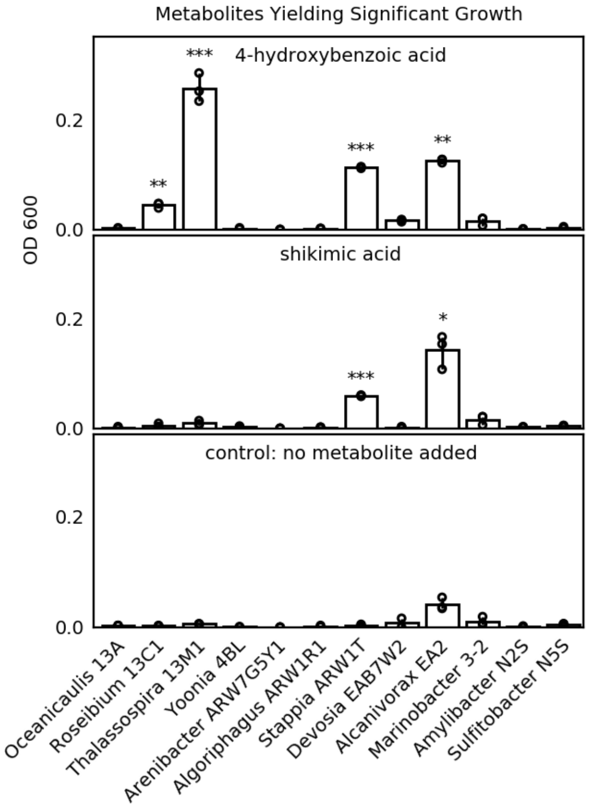

**Supplementary Figure S7.**

Responses of individual ASVs to exogenous addition of 4-hydroxybenzoic acid. All 31 ASVs with mean relative abundance above 0.1% are included. Horizontal bars and asterisks at the top indicate statistical significance. Significance codes: 0 '***' 0.001 '**' 0.01 '*' 0.05 '.' 0.1.

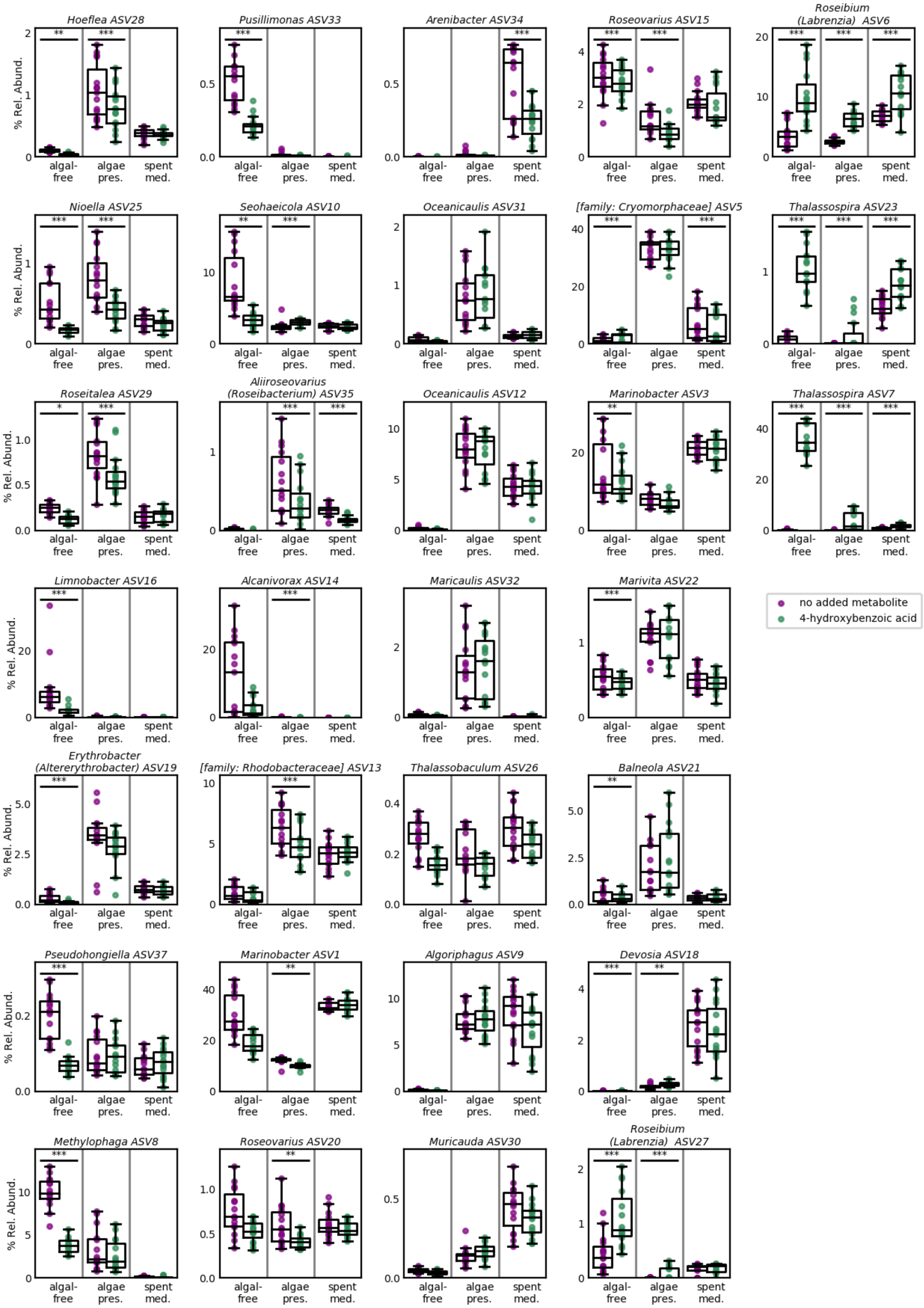
Hoeflea ASV28
Pusillimonas ASV33
Arenibacter ASV34
Roseovarius ASV15
Roseibium (Labrenzia) ASV6
Nioella ASV25
Seohaeicola ASV10
Oceanicaulis ASV31
[family: Cryomorphaceae] ASV5
Thalassospira ASV23
Roseitalea ASV29
Aliiroseovarius (Roseibacterium) ASV35
Oceanicaulis ASV12
Marinobacter ASV3
Thalassospira ASV7
Limnobacter ASV16
Alcanivorax ASV14
Maricaulis ASV32
Marivita ASV22
Erythrobacter (Altererythrobacter) ASV19
[family: Rhodobacteraceae] ASV13
Thalassobaculum ASV26
Balneola ASV21
Pseudohongiella ASV37
Marinobacter ASV1
Algoriphagus ASV9
Devosia ASV18
Methylophaga ASV8
Roseovarius ASV20
Muricauda ASV30
Roseibium (Labrenzia) ASV27
% Rel. Abund.
algal-free
algae pres.
spent med.
no added metabolite
4-hydroxybenzoic acid

**Supplementary Figure S8.**

Responses of individual ASVs to exogenous addition of lumichrome. All 31 ASVs with mean relative abundance above 0.1% are included. Horizontal bars and asterisks at the top indicate statistical significance. Significance codes: 0 ‘***’ 0.001 ‘**’ 0.01 ‘*’ 0.05 ‘.’ 0.1.

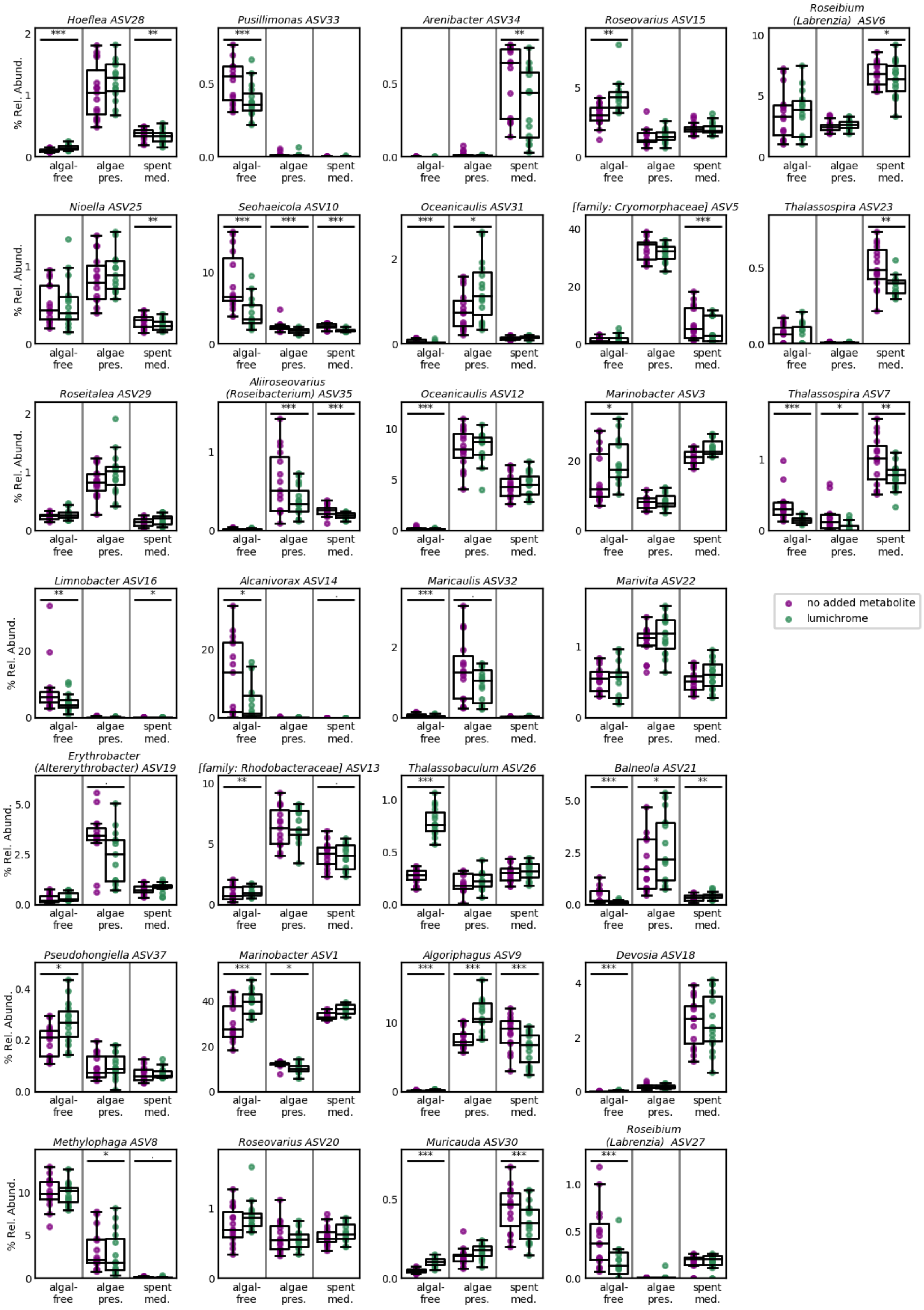
Hoeflea ASV28
Pusillimonas ASV33
Arenibacter ASV34
Roseovarius ASV15
Roseibium (Labrenzia) ASV6
Nioella ASV25
Seohaeicola ASV10
Oceanicaulis ASV31
[family: Cryomorphaceae] ASV5
Thalassospira ASV23
Roseitalea ASV29
Aliiroseovarius (Roseibacterium) ASV35
Oceanicaulis ASV12
Marinobacter ASV3
Thalassospira ASV7
Limnobacter ASV16
Alcanivorax ASV14
Maricaulis ASV32
Marivita ASV22
Erythrobacter (Altererythrobacter) ASV19
[family: Rhodobacteraceae] ASV13
Thalassobaculum ASV26
Balneola ASV21
Pseudohongiella ASV37
Marinobacter ASV1
Algoriphagus ASV9
Devosia ASV18
Methylophaga ASV8
Roseovarius ASV20
Muricauda ASV30
Roseibium (Labrenzia) ASV27
% Rel. Abund.
algal-free
algae pres.
spent med.
no added metabolite
lumichrome
